# Reprogramming of the intestinal epithelial-immune cell interactome during SARS-CoV-2 infection

**DOI:** 10.1101/2021.08.09.455656

**Authors:** Martina Poletti, Agatha Treveil, Luca Csabai, Leila Gul, Dezso Modos, Matthew Madgwick, Marton Olbei, Balazs Bohar, Alberto Valdeolivas, Denes Turei, Bram Verstockt, Sergio Triana, Theodore Alexandrov, Julio Saez-Rodriguez, Megan L. Stanifer, Steeve Boulant, Tamas Korcsmaros

## Abstract

Severe acute respiratory syndrome coronavirus 2 (SARS-CoV-2) represents an unprecedented worldwide health problem. Although the primary site of infection is the lung, growing evidence points towards a crucial role of the intestinal epithelium. Yet, the exact effects of viral infection and the role of intestinal epithelial-immune cell interactions in mediating the inflammatory response are not known. In this work, we apply network biology approaches to single-cell RNA-seq data from SARS-CoV-2 infected human ileal and colonic organoids to investigate how altered intracellular pathways upon infection in intestinal enterocytes leads to modified epithelial-immune crosstalk. We point out specific epithelial-immune interactions which could help SARS-CoV-2 evade the immune response. By integrating our data with existing experimental data, we provide a set of epithelial ligands likely to drive the inflammatory response upon infection. Our integrated analysis of intra- and inter-cellular molecular networks contribute to finding potential drug targets, and suggest using existing anti-inflammatory therapies in the gut as promising drug repurposing strategies against COVID-19.

## Introduction

Since the first reported case in the province of Wuhan (China), severe acute respiratory syndrome coronavirus 2 (SARS-CoV-2) has spread to almost every country in the world (Hui et al., 2020; Wu et al., 2020), posing an extraordinary threat to global public health (Deng and Peng, 2020; Han et al., 2020). Transmitted through respiratory droplets, aerosols and fomites, the virus can be detected in upper respiratory tract samples and is believed to primarily target airway and alveolar epithelial cells, vascular endothelial cells and lung-resident macrophages (Tay et al., 2020). Once inside the host cell, SARS-CoV-2 releases viral RNAs which can be translated into proteins using host machinery (Merino et al., 2021).

SARS-CoV-2 infection is not limited to the lungs: other organs can be infected too, including the heart, kidney, brain, and the intestine (Gupta et al., 2020). In addition to directly infecting key organs, the main hurdle of SARS-CoV-2 infection is that, in some severe cases, it generates an excessive inflammatory response mediated by both the innate and adaptive immune systems (Olbei et al., 2021). The overactivated inflammatory response, also known as cytokine release syndrome (CRS) or cytokine storm, is the result of high levels of circulating cytokines and chemokines, and it is thought to be responsible for the severe COVID symptoms some patients experience (Arunachalam et al., 2020). Yet, there is no clear understanding of which particular inflammatory pathways and cell types are driving these damaging inflammatory responses, and whether some organs are more important than others in initiating this (Stone et al., 2020). The role of gut microbes and previous infections were also raised as potential risk-increasing factors (Földvári-Nagy et al., 2021).

COVID-19 patients with severe symptoms show elevated expression of inflammatory cytokines (IL-2, IL-4, IL-6, IL-10 and IL-18; (Gu et al., 2020; Park et al., 2021) that are correlated with elevated levels of the gut inflammatory marker *faecal calprotectin* and an altered microbiome (Effenberger et al., 2020; Zuo et al., 2020). COVID-19 patients also often present various gastrointestinal (GI) symptoms such as vomiting, diarrhoea and abdominal pain (Chen et al., 2020; Guo et al., 2020; Lin et al., 2020). Interestingly, patients with GI symptoms show decreased production of key proinflammatory cytokines and reduced disease severity and mortality following SARS-CoV-2 infection, indicating a potential role of the gut in the disease course (Livanos et al., 2021).

Recently, human intestinal organoids have been used as a tool to study SARS-CoV-2 infection in the gut and the inflammatory responses of specific intestinal epithelial cell types (Lamers et al., 2020; Stanifer et al., 2020; Triana et al., 2021a; Zang et al., 2020). These studies provided evidence that SARS-CoV-2 is able to infect and actively replicate in human intestinal cells (Lamers et al., 2020). From the organoid experiments, we learned that while most intestinal subpopulations are susceptible to SARS-CoV-2, enterocytes are the most affected (Lamers et al., 2020; Triana et al., 2021a). Studies using human intestinal organoids also revealed that, contrary to the limited type I and type III interferon (IFN) immune response observed in the lungs (Blanco-Melo et al., 2020; Hadjadj et al., 2020), the response to SARS-CoV-2 infection in the gut is characterised by the production of IFN and interferon stimulated genes (ISGs), and found that IFNs may actually provide protection to the intestinal epithelial cells against SARS-CoV-2 (Stanifer et al., 2020).

In inflammatory bowel disease (IBD), intestinal inflammation is known to drive dysregulated epithelial-immune cell interactions, often manifesting in extra-intestinal diseases (Weidinger et al., 2021). Hence, the question arises as to whether COVID-19 patients may share a similar dysregulated inflammatory response driven by gut epithelial-immune interactions as that observed in IBD patients. Indeed, examination of human intestinal samples has shown infiltration of lymphocytes and other inflammatory mediators in the lamina propria upon SARS-CoV-2 infection, suggesting that infection of gut epithelial cells results in the activation of local immune populations (Guo et al., 2021). Yet, the exact effects of viral infection in the gut and the role of epithelial cell–immune cell interaction in mediating the inflammatory response of the body are not known.

To our knowledge, no study has been carried out so far to analyze epithelial–immune crosstalk in the gastrointestinal tract upon SARS-CoV-2 infection. Hence, in this study we aim to model the effect of viral infection in host intestinal cells, the role of intestinal epithelial cell–immune cell crosstalk during infection, as well as their contribution to the inflammatory response. As miRNA-like sequences were recently found in the SARS-CoV-2 genome, and their potential role and targets predicted during infection (Mirzaei et al., 2021; Saçar Demirci and Adan, 2020), in a separate analysis, we assess the potential role of such viral regulators. Furthermore, we aim to assess potential tissue-specific differences between colon and ileum in these effects. To do this, we used previously generated single-cell RNA sequencing (scRNA-seq) data of SARS-CoV-2 infected ileum and colon-derived human organoids (Triana et al., 2021a) and of gut-resident immune cell populations (Martin et al., 2019; Smillie et al., 2019), as well as SARS-CoV-2–human miRNA/protein–protein interactions. We employ two independent tools, ViralLink and CARNIVAL, to reconstruct intracellular and intercellular networks, connecting intestinal epithelial cells and resident immune cells upon infection. With our integrated analysis, we provide a better understanding of the effect of viral infection on intestinal epithelial cells, and the role of intestinal epithelial– immune cell crosstalk during SARS-CoV-2 infection. Ultimately, our analyses may help to find key intercellular inflammatory pathways involved in these crosstalks, which could pave the way for potential successful strategies against the cytokine release syndrome associated-symptoms observed in severe cases of COVID-19.

## Methods

### Intercellular analysis

#### Input data

##### Intestinal epithelial cells

Single cell transcriptomics data of colonoids and enteroids infected with SARS-CoV-2 was obtained from (Triana et al., 2021b) The R packages ‘Mast’ and ‘Seurat’ were used to identify differentially expressed genes upon infection with SARS-CoV-2 for each epithelial cell type. Specifically, directly infected or bystander cells from organoids treated with SARS-CoV-2 for 24 hours were compared with the equivalent cell type from uninfected organoids. Any genes with adjusted p value ≤ 0.05 and |log2 fold change (FC)| ≥ 0.5 were considered significantly differentially expressed. Differential expression could only be calculated for cell types within a condition where data was available from ≥ 3 individual cells.

##### Intestinal resident immune cells

Single cell expression data from ileal and colonic resident immune cells was obtained from (Martin et al., 2019) and (Smillie et al., 2019), respectively. For the analyses, data from healthy samples and uninflamed Crohn’s disease samples was used for colonic and ileal immune cell populations, respectively. Following removal of all genes with count = 0, normalised log2 counts across all samples (separately for each cell type) were fitted to a gaussian kernel (Beal, 2017). All genes with expression values above mean minus three standard deviations were considered as expressed genes for the given cell type in the given intestinal location. For the intercellular ligand-receptor predictions, a representative collection of immune cells relevant in gut inflammation and SARS-CoV-2 infection based on previous literature was selected, which included all macrophages, T cells, B cells, plasma cells, ILCs, mast cells and a representative group of dendritic cells (Filbin et al., 2020; Martin et al., 2019; Schultze and Aschenbrenner, 2021; Sette and Crotty, 2021; Smillie et al., 2019). Cell type labels were maintained as originally published.

#### Defining ligand-receptor interactions between cell types

A list of ligand-receptor interactions was obtained from OmniPath on 23 September 2020 using the ‘OmnipathR’ R package (Türei et al., 2021). Source databases used to retrieve the ligand-receptor interactions through OmnipathR included six independent resources (CellPhoneDB, HPMR, Ramilowski 2015, Guide2Pharma, Kirouac 2010, Gene Ontology) (Ashburner et al., 2000; Ben-Shlomo et al., 2003; Kirouac et al., 2010; Pawson et al., 2014; Ramilowski et al., 2015; Vento-Tormo et al., 2018). No weighing was performed on ligand-receptor interactions, and protein complexes were dealt with by including each of their individual proteins in the list.

Ligand-receptor interactions (intercellular interactions; full list available at https://github.com/korcsmarosgroup/gut-COVID) were predicted between epithelial cells types and resident immune cells according to the following conditions:

1. The ligand is differentially expressed in the epithelial cell (upon SARS-CoV-2 infection — in directly infected or bystander cells)
2. The receptor is expressed in the immune cell in the relevant dataset (ie, ileal or colonic immune cells)
3. The ligand-receptor interaction is present in OmniPath

Intercellular interactions were defined separately for directly infected epithelial cells and bystander epithelial cell populations in the ileum and in the colon. Enteroid epithelial data was paired with ileal immune cell data (Martin et al., 2019), while colonoid epithelial data was paired with colonic immune cell data (Smillie et al., 2019). Intercellular interactions were defined between every possible pair of epithelial cells and immune cells for each condition. Interactions derived from upregulated ligands (“upregulated interactions”) were evaluated separately from interactions derived from downregulated ligands (“downregulated interactions”).

#### Scoring of ligands, receptors and immune cell types involved in ligand-receptor interactions

To assess the importance of specific ligands, receptors and immune cell types, additional parameters were computed using the ligand-receptor network. First, the number of interactions between each epithelial and immune cell type was computed by summing up all the possible interactions between each differentially expressed epithelial ligand and each of the receptors expressed by the specific immune cell type. Second, the number of immune cell types involved in each ligand-receptor pair was computed by counting the number of different immune cell types which were expressing the receiving receptor. Third, for each ligand, a “sum of receptor expression value” was computed for each interacting immune cell type, based on the number of interacting receptors and the mean expression level of the interacting receptors.

#### Data visualisation

Venn diagrams were generated using the ‘gplots’ R package. Heatmaps were generated using the ‘ggplot2’ and ‘pheatmap’ packages. Barplots were generated with the ‘ggplot2’ package. Network visualisations were done using Cytoscape (version 3.8.2) (Shannon et al. 2003; Su et al. 2014). All scripts used to generate these plots are available on the Github repository of the project (https://github.com/korcsmarosgroup/gut-COVID).

### Intracellular analysis

Two previously established tools were employed to predict the effect of SARS-CoV-2 infection on intestinal epithelial cells: ViralLink and CARNIVAL (Liu et al., 2019; Treveil et al., 2021). Both tools, using related but distinct methods, infer causal molecular interaction networks. These networks link perturbed human proteins predicted to interact with SARS-CoV-2 viral proteins or miRNAs, to transcription factors known to regulate the observed differentially expressed ligands in infected epithelial cells.

#### Input data

To reconstruct the intracellular causal networks, three different *a priori* interactions datasets were used. Information on viral proteins and their interacting human binding partners was obtained from the SARS-CoV-2 collection of the IntAct database on 1st October 2020 (Hermjakob et al., 2004; Orchard et al., 2014). Predicted SARS-CoV-2 miRNAs and their putative human binding partners were obtained from (Saçar Demirci and Adan, 2020). Intermediary signalling protein interactions known to occur in humans were obtained from the core protein-protein interaction (PPI) layer of the OmniPath collection using the ‘OmnipathR’ R package on 7th October 2020 (Türei et al., 2016). Only directed and signed interactions were included. Interactions between human transcription factors (TFs) and their target genes (TG) were obtained from the DoRothEA collection using the DoRothEA R package on 7th October 2020 (Garcia-Alonso et al., 2019). Only signed interactions of the top three confidence levels (A, B, C) were included.

Normalised transcript counts and differentially expressed ligands were obtained from single cell transcriptomics data of colonoids and enteroids infected with SARS-CoV-2 obtained from (Triana et al., 2021a) as previously described.

#### ViralLink pipeline

Intracellular causal networks were inferred using the ViralLink pipeline, as described in (Treveil et al., 2021). Briefly, a list of expressed genes in infected immature enterocytes (originally known as “immature enterocytes 2” (MMP7+, MUC1+, CXCL1+)) from SARS-CoV-2-infected ileal and colonic organoids (Triana et al., 2021a) was generated from a normalised count table by fitting a gaussian kernel (Beal, 2017). The list of expressed genes in the infected immature enterocytes population was subsequently used to filter the *a priori* molecular interactions from OmniPath and DoRothEA, to obtain PPI and TF-TG sub-networks where both interacting molecules are expressed (described as “contextualised” networks). Transcription factors regulating the differentially expressed ligands were predicted using the contextualised DoRothEA TF—TG interactions and scored as described in (Treveil et al., 2021). Human binding proteins of viral proteins and viral miRNAs obtained from the IntAct database (Hermjakob et al., 2004; Orchard et al., 2014) and (Saçar Demirci and Adan, 2020), respectively, were connected to the listed TFs through the contextualised PPIs using a network diffusion approach called Tied Diffusion Through Interacting Events (TieDIE) (Paull et al., 2013). In this model, all viral protein—human binding protein interactions were assumed to be inhibitory in action, based on likely biological function, and given a lack of clear literature evidence of proven action. All viral miRNA—human binding protein interactions were set as inhibitory based on biological action of miRNAs (Huang et al., 2011). The final reconstructed network contains “nodes”, which refers to the interacting partners, and “edges”, which refers to the interaction between nodes. Nodes include viral proteins and miRNAs, human binding proteins, intermediary signalling proteins, TFs and differentially expressed ligands. Edges include activatory or inhibitory interactions.

For both ileal and colonic data, separate networks were generated using the viral miRNA and viral protein as perturbations, and subsequently joined using the “Merge” function within Cytoscape to generate the final intracellular network. Nodes and edges were annotated according to their involvement in networks downstream of viral miRNAs or proteins. Further analyses were performed separately on the different layers of the network: miRNA specific, protein specific or shared nodes.

#### CARNIVAL pipeline

Intracellular causal networks were inferred using CARNIVAL and associated tools for analyses of expression data as described in (Liu et al., 2019). For simplicity, we refer to the pipeline as described in (Liu et al., 2019) as the CARNIVAL pipeline. Briefly, PROGENy is used to infer pathway activity from the log2 FC of the infected immature enterocytes 2 gene expression data (Schubert et al., 2018). Next, using the TF-TGs interactions (from DoRothEA) and the differential expression data from infected organoids, VIPER was used to score TF activity based on enriched regulon analysis (Alvarez et al., 2016). Here, only the top 25 TFs regulating at least 15 target genes were taken forward, and a correction for pleiotropic regulation was included. Finally, CARNIVAL applied an integer linear programming approach to identify the most likely paths between human interaction partners of viral proteins or miRNAs and the selected TFs, through PPIs from OmniPath, considering the activity scores from PROGENy and VIPER. Viral protein—human binding protein interactions signs were specified to CARNIVAL as ‘inhibitory’, based on likely biological function, and given a lack of clear literature evidence of proven action. All viral miRNA— human binding protein interactions were also set as inhibitory based on biological action of miRNAs (Huang et al., 2011).

### Network functional analysis

Functional overrepresentation analysis was performed on the networks constructed as above-mentioned using the R packages ‘ClusterProfiler’ and ‘ReactomePA’, for Gene Ontology (GO) (Ashburner et al., 2000)) and for Reactome (Fabregat et al., 2018; Yu and He, 2016; Yu et al., 2012) annotations, respectively. For the intercellular network, the analysis was carried out separately for ligand-receptor intercellular interactions driven by either upregulated or downregulated ligands. A complete list of ligand-receptor interactions is available in the GitHub repository of the project (https://github.com/korcsmarosgroup/gut-COVID). For the upregulated interactions, a list of upregulated ligands and their connecting immune receptors was used as the test. For the downregulated interactions, a list of downregulated ligands and their connecting immune receptors was used. Where a list of ligands plus receptors contained <5 genes, it was excluded from the analysis. All ligands and receptors from the original ligand-receptor network used as prior knowledge input for the intercellular analysis was used as the statistical background.

For the intracellular network, the analysis has been done separately for each of the sub-networks (viral protein specific, viral miRNA specific, or shared). For each sub-network, a set of genes that were human binding proteins, intermediary proteins and TFs in the network (“PPI layer”) was used as a test list, and a set of all nodes from the original OmniPath PPI interaction network used as prior knowledge input for the intracellular analysis was used as the statistical background. For the Reactome pathway enrichment analysis the IDs were converted to Entrez Gene ID within the ‘ReactomePA’ package. Functional categories with adjusted p value ≤ 0.05 and with gene count > 3 were considered significantly overrepresented.

## Selection of ligands involved in the inflammatory process

To assess the importance of specific ligands in driving the inflammatory process upon SARS-CoV-2 infection, the list of differentially expressed ligands in infected immature enterocytes in both colon and ileum was validated using independent data from three previously published studies. To identify ligands whose expression was induced by cytokines, ligands were compared to DEGs in human colonic organoids exposed to cytokines from (Pavlidis et al., 2021). To identify ligands already known to influence immune cell population, ligands were compared to two databases: ImmunoGlobe, a manually curated intercellular immune interaction network (Atallah et al., 2020), and ImmunoeXpresso, a collection of cell–cytokine interactions generated through text mining (Kveler et al., 2018). Finally, to identify ligands that could directly explain blood cytokine level changes in COVID-19 patients via direct immune cell regulation, ligands were compared to the data from a large dataset we recently manually compiled using COVID-19 patient publications (Olbei et al., 2021).

## Data availability

The workflow (and necessary input data) and the full ligand-receptor interaction tables are available in the GitHub repository of the project (https://github.com/korcsmarosgroup/gut-COVID). All other relevant data is in the main text and in supplementary files.

## Results

### Reconstructing an epithelial-immune interactome

Our previously published data on ileal and colonic human organoids infected with SARS-CoV-2 suggested that immature enterocytes were the main epithelial population affected by infection (Triana et al., 2021a). In this study, we wanted to further investigate the effects of epithelial infection on epithelial-immune cell crosstalk in the gut by integrating single cell data and network biology approaches (**Figure 1**).

**Figure 1.**
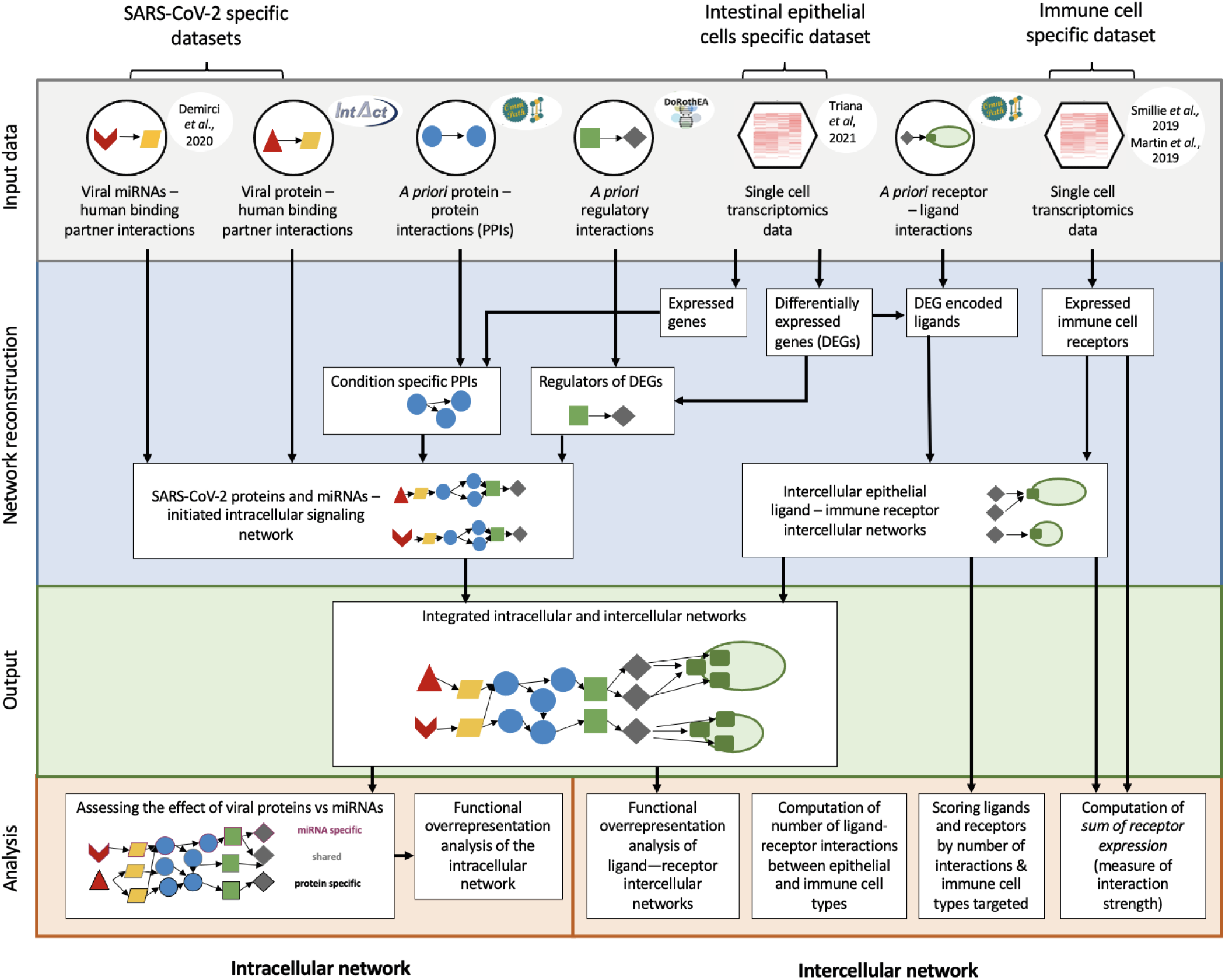
Integrated workflow to analyse the intracellular and intercellular effect of SARS-CoV-2 in the gut. Schematic workflow illustrating the different analytical steps used to construct the intracellular and intercellular signalling networks between epithelial cells in SARS-CoV-2 infected intestinal organoids (ileal and colonic organoids, 24 hrs infection) and immune cell types.

To do this, we integrated epithelial cell (Triana et al., 2021a) and immune cell (Martin et al., 2019; Smillie et al., 2019) RNAseq data and *a priori* knowledge on ligand-receptor interactions (Türei et al., 2021) to construct intercellular networks connecting epithelial cells and resident immune cells (**Figure 1**). Final constructed intercellular networks consisted of differentially expressed epithelial cell ligands binding to receptors expressed on healthy immune cells (**Figure 1**). In accordance with our previous findings (Triana et al., 2021a), immature enterocytes (originally known as “immature enterocytes 2”, an enterocyte subpopulation characterized by MMP7+, MUC1+, CXCL1+) were the epithelial population whose ligands were most affected upon infection, among the different cell types studied **(Supplementary Figure 1)**. Additionally, directly infected cells, compared to bystander cells, showed the highest number of differentially expressed ligands in both colon and ileum (**Supplementary Figure 1**).

To identify the effect of epithelial infection on epithelial-immune cell crosstalk, we looked at the putative number of ligand-receptor interactions between each epithelial cell and immune cell types (**Figure 1**). Here, for each epithelial-immune cell type pair, the number of potential interactions was computed by summing up all the possible interactions between each set of up or downregulated epithelial ligands and each of the receptors expressed by the specific immune cell type (from (Martin et al., 2019) and (Smillie et al., 2019)) (**Figure 2A**). Both bystander and infected cell populations were affected by viral infection, but directly infected cell populations had a higher number of predicted interactions with immune cells than bystander cell populations in both ileum and colon, supporting a role for direct viral infection in altering intercellular signalling in the gut (**Figure 2A**). In the colon, the higher number of epithelial-immune interactions involved downregulated ligands, with plasma cells being the main immune cell type involved in these interactions, and CD4+/CD8+ T cells, macrophages and dendritic cells (DCs) to a lesser extent (**Figure 2A**). Conversely, in the ileum, the highest number of interactions involved upregulated ligands, with IgA plasma cells, T resident memory (Trm) cells, dendritic cells and resident macrophages as the main immune cell types involved in these interactions (**Figure 2A**). Notably, the higher number of interactions in the ileum was not a result of a higher number of upregulated ligands (20), as this was similar to the number of downregulated ones (24) (**Figure 2A, Supplementary Figure 1**). Instead, the higher number of interactions was driven by upregulated ligands binding to multiple receptors on each immune cell targeted (not shown).

**Figure 2.**
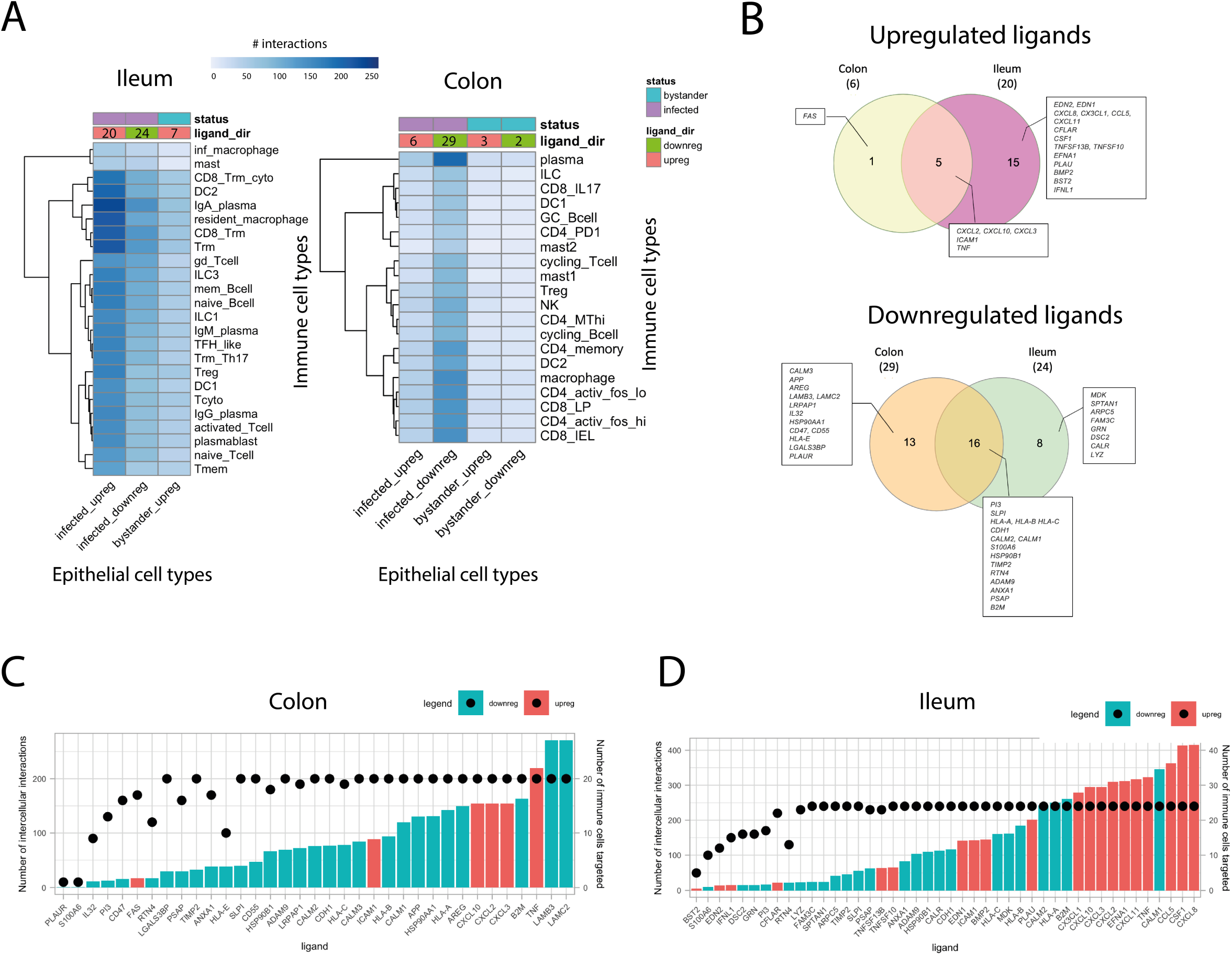
Differentially expressed ligands driving upregulated and downregulated intercellular interactions between colonic and ileal infected immature enterocytes and resident immune cells upon infection in the colon and ileum. **(A)** Heatplot showing the number of interactions between immature enterocytes and resident immune cells. Interactions driven by upregulated and downregulated ligands (ligand_dir) are shown separately for infected and bystander cells (status), and for ileum and colonic organoids. The intensity of the color indicates the number of immune cell types whose receptor is targeted by the epithelial cells ligands. The numbers on the ligand_dir row refer to the number of upregulated or downregulated ligands driving the indicated interactions with immune cells for the different groups/conditions. Abbreviations: *<u>Ileum</u>*: inf_macrophage, infected macrophage; mast, mast cell, CD8_Trm_cyto, Resident memory cytotoxic T cell; DC2, dendritic cell 2; Trm, Tissue-resident memory T cell, gd_Tcell, Gamma delta (γδ) T cells; ILC, Innate lymphoid cell; mem_Bcell, memory B cell; naive_Bcell, naive B cell; TFH_like, T follicular helper cells; Trm_Th17, Tissue-resident memory Th17 cells; Treg, Regulatory T cell; Tcyto, Cytotoxic T cell; Tmem, Memory T cells. *<u>Colon</u>*: ILC, Innate lymphoid cell; CD8_IL17, IL-17+ CD8+ T cells; DC, dendritic cells; GC_Bcell, Germinal center B cells; CD4_PD1, mast, mast cell; Treg, Regulatory T cell; NK, Natural Killer cell, CD4_MThi, high mitochondrial CD4+ T cell; CD4_memory, CD4+ Memory T cell, CD4_activ_fos_high, activated CD4+ T cells (high/low c-fos); CD8_LP, CD8+ lymphocyte-predominant cells, CD8_IEL, CD8+ intraepithelial lymphocytes. **(B)** Venn diagrams showing the number of ligands of the infected immature enterocytes - immune cells intercellular network that are unique or shared between the ileum and colon. Upregulated and downregulated ligands are shown separately. **(C, D)** Bar plot showing the upregulated and downregulated ligands in the colonic (C) and ileal (D) infected immature enterocytes - immune cell network scored by number of interactions (height of the bar plot) and number of immune cells targeted (black dots). Upregulated ligands are shown in red and downregulated ligands in blue.

### The infected epithelial signalling network drives the epithelial-immune interactome

To further understand how SARS-CoV-2 infection results in altered ligand expression, we investigated the effect of SARS-CoV-2 infection on intracellular signalling in directly infected immature enterocytes. Using two independent bioinformatics tools, ViralLink and CARNIVAL, we constructed an intracellular causal network linking perturbed human proteins interacting with SARS-CoV-2 viral proteins or miRNAs to activated transcription factors (TFs) regulating the differentially expressed ligands upon infection, through altered intracellular protein-protein signalling cascades (**Figure 1** **and Methods**). To spot tissue-specific differences in infection response, two separate causal networks were constructed for infected immature enterocytes of the ileum and colon (**Figure 1** **and Methods**). Furthermore, to assess the contribution of viral proteins or miRNAs in altering the intracellular signalling cascade, separate layers of the networks were built distinguishing altered signalling stemming from upstream perturbations caused by SARS-CoV-2 miRNAs, proteins or both (**Figure 1** **and Methods**).

Colonic and ileal intracellular networks generated using ViralLink were similar in terms of size and network characteristics for ileum and colon, when considering the diameter, characteristic path length, average number of neighbours, and number of molecular entities (nodes, miRNAs, genes or proteins) and molecular interactions (edges, activatory or inhibitory) **(Supplementary Figure 2)**. A complete description of network characteristics for both colonic and ileal networks is available in the **Supplementary Information**. Notably, upstream signalling was predicted for 22 out of the initial 35 differentially expressed ligands (29 down- and 6 up-regulated) in the colon, and for 28 out of 44 differentially expressed ligands (24 down- and 20 up-regulated) for the ileum (**Supplementary Figure 2, 3 & 4**). These numbers are lower than those predicted to be differentially expressed upon infection by (Triana et al., 2021a), indicating that some ligands are not affected by direct upstream signalling changes but by more complex mechanisms, or the original knowledge network used as input for the analysis did not contain information about such ligands (Menche et al., 2015) **(Supplementary Figure 1 & 2).**

To understand how SARS-CoV-2 infection in immature enterocytes affected their function through the modulation of intracellular signalling, we performed a functional overrepresentation analysis (Gene Ontology (GO) and Reactome) of the protein-protein interaction (“PPI”) layer (Ashburner et al., 2000; Fabregat et al., 2018; Yu and He, 2016; Yu et al., 2012) (**Figure 1** **and Methods**). This analysis was performed separately for each sub-network (viral proteins, viral miRNA, both) to assess the contributions of SARS-CoV-2 miRNAs or proteins to the changes observed (**Figure 1** **and Methods**). Functional analysis revealed an overrepresentation of pathways related to inflammation and chemotaxis (NF-kB signalling, interleukin signalling, chemokine signalling) in both ileum and colon (**Figure 3A, 4B**). Additionally, we found the overrepresentation of functions related to interferon signalling and MAPK signalling being overrepresented uniquely in the ileum in both viral protein and miRNA intracellular networks (**Figure 3B**). An overrepresentation of laminin-driven interaction pathways, which we observed uniquely for viral miRNA intracellular network in both ileum and colon, could be indicative of an increased recruitment and adhesion of immune cells following infection (**Figure 3A, 3B**). Furthermore, we found an overrepresentation of pathways related to negative regulation of apoptosis, cell cycle, cell proliferation and growth in both ileum and colon, suggesting an effect of SARS-CoV-2 on epithelial cell tissue renewal (**Figure 3A, 3B**). Interestingly, uniquely in the viral-protein subnetwork, we found an overrepresentation of WNT signalling, which is involved in stem cell renewal, in both ileum and colon, as well as pathways related to the establishment of cell and tissue polarity uniquely in the colon, which could indicate an attempt for tissue healing following viral infection (**Figure 3A, 3B**).

**Figure 3.**
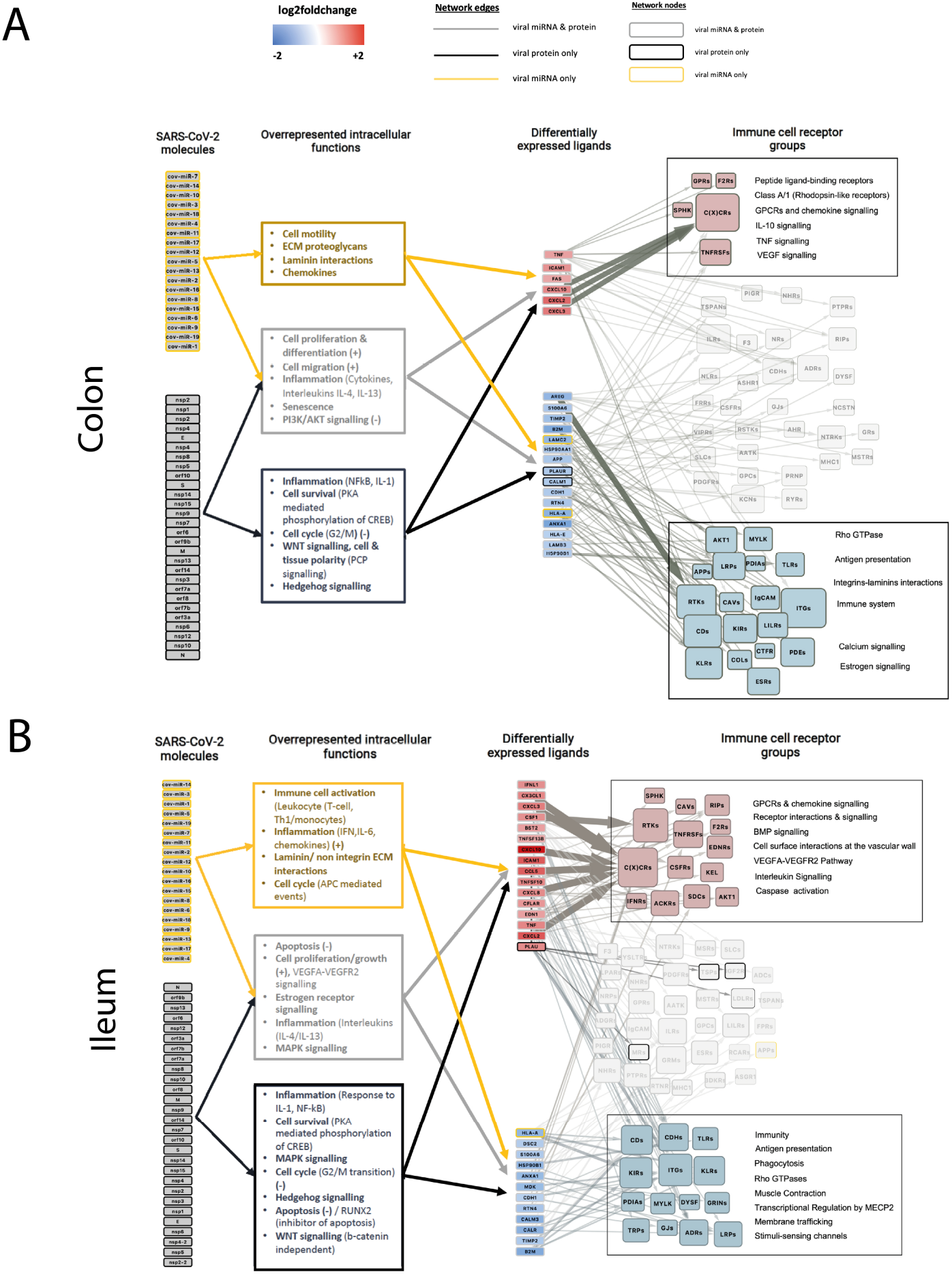
Overview of intracellular and intercellular signalling of ileal and colonic infected immature enterocytes upon SARS-CoV-2 infection. **(A, B)** Overview of intracellular and intercellular signalling upon SARS-CoV-2 infection in colonic (A) and ileal (B) infected immature enterocytes and immune cell populations. From left to right: signalling cascade going from SARS-CoV-2 molecules (proteins or miRNAs) to differentially expressed ligands on immature enterocytes and binding receptor groups on immune cells. <u>Intracellular network:</u> SARS-COV-2 molecules are grouped separately if they are viral proteins (bottom) or miRNAs (top). Differentially expressed ligands for which no upstream signalling was identified, but downstream intercellular connections were predicted are excluded from this figure. Differentially expressed ligands are grouped based on the direction of regulation, which is indicated with blue when downregulated (bottom) and red when upregulated (top) when comparing SARS-CoV-2 infected vs uninfected conditions. Colors of the nodes and of the functional analysis indicate if the original network was a miRNA only (yellow), viral protein only (black) or both viral protein and miRNA (grey). Functional overrepresentation analysis was carried out for the “PPI layer” of the intracellular network which includes human binding proteins, intermediary signalling proteins and TFs (adj p value < 0.05, n > 3). <u>Intercellular network</u>: Size of the receptor node represents the sum of receptors within the group targeted by each incoming ligand. Functional analysis is indicated for ligand-receptor groups. Receptor groups layout is based on whether they contributed to the functional analysis of upregulated interactions (red) or downregulated interactions (blue). Receptor groups not contributing to any functions are indicated in light grey.

**Figure 4.**
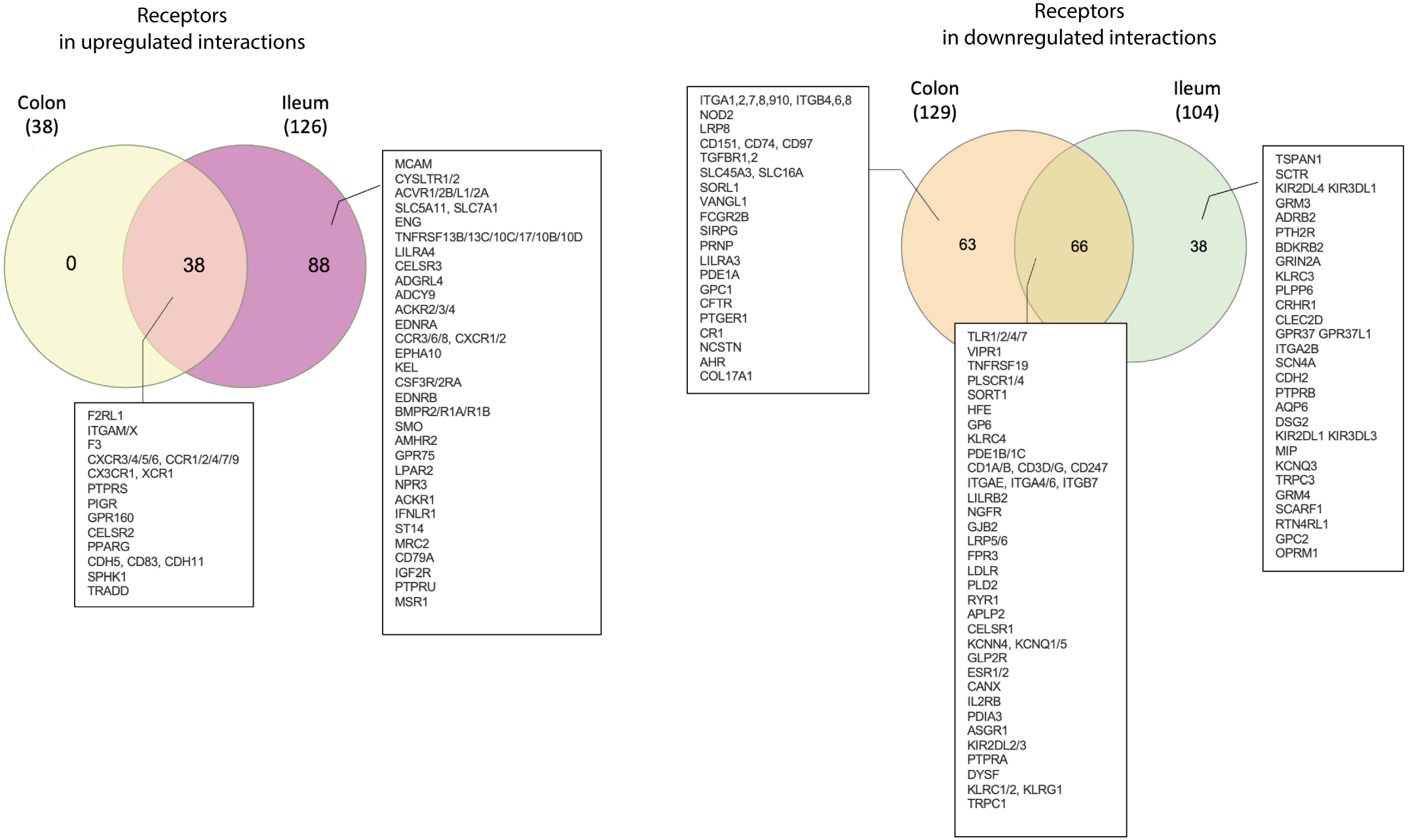
Overview of receptors involved in upregulated and downregulated interactions between infected immature enterocytes and resident immune cells upon infection in the colon and ileum. Venn diagrams showing the number of receptors in the infected immature enterocytes - immune cells intercellular networks that are unique or shared between the ileal and colonic networks. Receptors targeted by upregulated ligands and downregulated ligands are shown separately.

Intracellular signalling networks built with CARNIVAL as an independent tool were of much smaller sizes compared to those built with ViralLink (**Supplementary Figure 2**). Compared to networks built with ViralLink where all possible interaction paths are explored, CARNIVAL networks represent the most optimal paths based on the given input constraints, hence they are very useful to understand specific molecular mechanisms and modulators upon SARS-CoV-2 infection. Functional overrepresentation analysis of the PPI layer of the CARNIVAL networks confirmed similar modulated functions upon infection compared to those found in the ViralLink networks **(Supplementary Figure 3 & 4)**. Using these networks, we predicted which transcription factors regulated upstream the altered ligand expression upon infection in directly infected immature enterocytes. Both tissues shared similar transcription factors, including ATF2/3, FOS, JUN, STAT1, and NFKB1, which were all upregulated in both tissues (**Supplementary Figure 3 & 4**). These transcription factors play a role in interferon response (STAT1) and inflammation (NFKB1), anti-apoptosis and cell growth (ATF2/3), cell proliferation and differentiation (JUN, FOS), suggesting an increase in these these functions upon SARS-CoV-2 infection in both colon and ileum. Interestingly, viral miRNAs that were predicted to regulate upstream the altered intracellular signalling were mostly different between colon and ileum (miR_10,11,16,18 in the colon and miR_4,5,6,18 in the ileum). Additionally, by analysing these networks, we observed that NOTCH1 and SMAD4, seem to be central to the intracellular signalling cascade in the colon, by receiving several signals driven by viral miRNAs and viral proteins, respectively (**Supplementary Figure 3**). Interestingly, both the Notch and TGF-β SMAD-dependant signaling pathways are involved in intestinal epithelial cell homeostasis, including stem cell maintenance, progenitor cell proliferation (Carulli et al., 2015) and maintenance of cell differentiation (Yamada et al., 2013), suggesting a modulation of these pathways upon infection. In the ileal network, JAK2 and CREB1, as well as SMAD2, SMAD 3 and ERK2 (MAPK1) seem to play a central role in the intracellular PPI signalling driven by viral miRNAs and viral proteins, respectively, and *JAK2* and both *SMAD2* and *SMAD3* were also upregulated upon infection (**Supplementary Figure 4**).

### Upregulated epithelial ligands upon infection impact pro-inflammatory responses and immune cell recruitment to the infected epithelium

To understand the functional impact of epithelial infection on intercellular communication, we looked at the ligand-receptor interactions driven by up and downregulated epithelial ligands in infected immature enterocytes upon infection in the colon and ileum (**Figure 1, Supplementary Figure 8, and Methods**). For each set of up and downregulated intercellular interactions, we looked at which ligands, receptors and immune cell types were involved in these intercellular interactions, assessing any potential similarities or differences between the colon and ileum (**Figure 1** **and Methods**).

Upregulated ligands in infected immature enterocytes were largely shared between colon and ileum, with one ligand (*FAS*) uniquely upregulated in the colon (**Figure 2B**). Shared upregulated ligands included mainly cytokines and chemokines (CXCL2/3/10 and tumor necrosis factor (TNF-a)) and the adhesion factor ICAM1. Interestingly, several additional chemokines (CSF1, CXCLs, TNFSFs) and adhesion factors (PLAU, EFNA) were upregulated in the ileum upon infection, which we did not find in the colon (**Figure 2B**). Additionally, 38 receptors on immune cells targeted by upregulated ligands in the colon were all shared with the ileum, and were mainly represented by chemokine receptors (CXCRs, CCRs) (**Figure 4**). Epithelial-immune interactions driven by upregulated ligands were also mostly shared in the colon and ileum (1 unique to colon, 219 unique to ileum, 66 shared) (**Figure 6** **& Supplementary Figure 8A, 8B)**.

Next, to understand which ligands were driving the most interactions with specific immune cell types, we scored them based on the number of ligand-receptor interactions they had with the different immune cell types analysed (**Figure 1** **and Methods).** Chemokines (CXCLs) and tumor necrosis factor alpha (TNF-a) were among epithelial ligands (**Figure 2C, 3D**), and chemokine receptors (CXCR 3,4,5,6 and CCR 1,2,5,7,9,10) among the receptors on immune cells driving the highest numbers of upregulated interactions in both tissues (**Figure 5A, 5B & Supplementary Figure 6A, 6B**), overall pointing towards an increased immune cell recruitment upon infection. The high number of upregulated interactions driven by chemokines could be attributable to the widespread presence of several different chemokine receptors on immune cells (**Supplementary Figure 8A, 8B**). Additionally, we found ileal-specific upregulated interactions involving Plasminogen Activator (PLAU), Ephrin A1 (EFNA1) and colony stimulating factor 1 (CSF1) binding to various receptors on immune cells, pointing towards an increased immune cell recruitment and adhesion **(Figure 6, Supplementary Figure 8B).** Finally, we found one colon-specific upregulated interaction between epithelial Fas Cell Surface Death Receptor (FAS) binding to receptor-interacting serine/threonine-protein kinase 1 (RIPK), pointing towards increased cell death upon infection (**Figure 6, Supplementary Figure 8A**).

**Figure 5.**
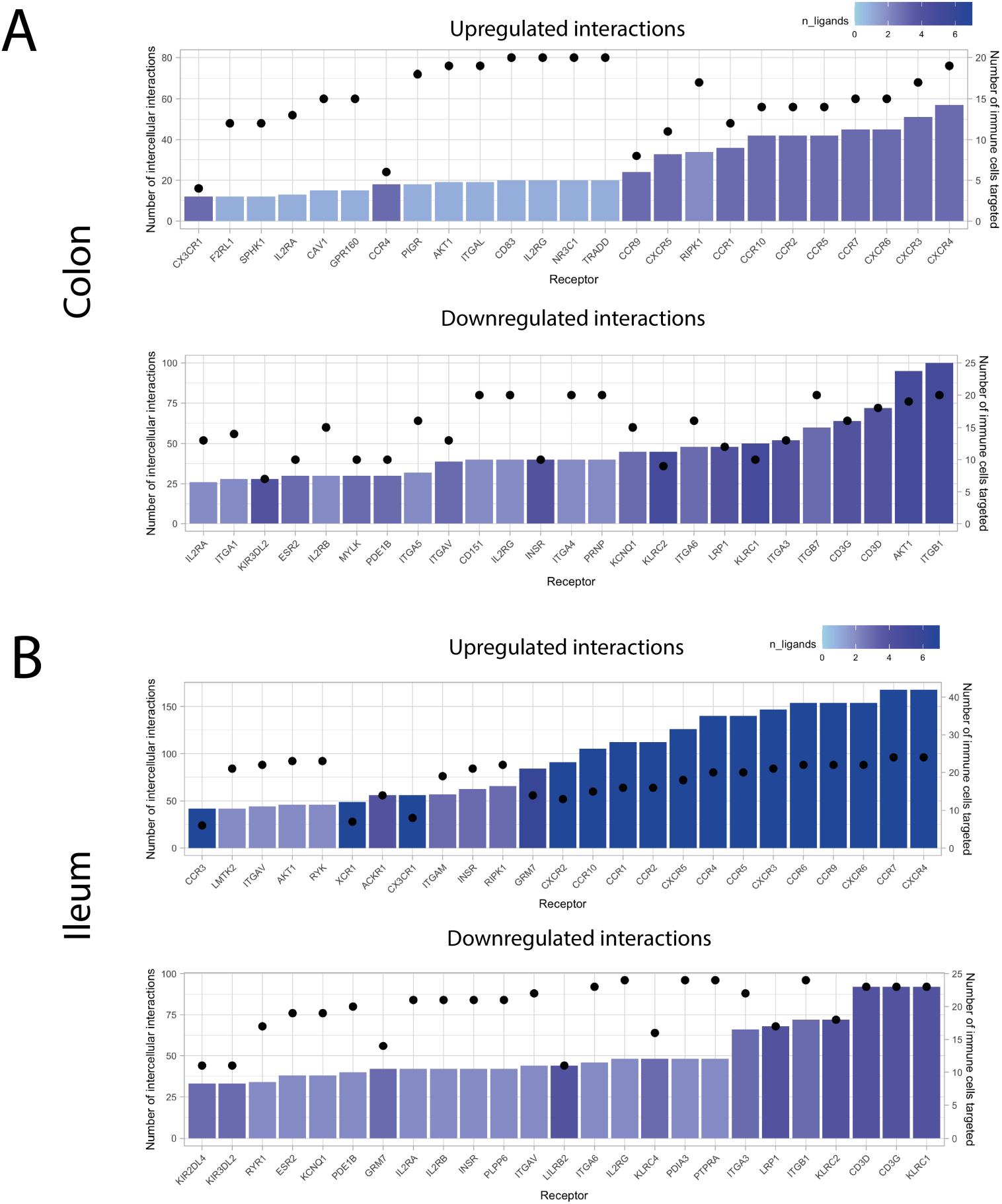
Receptors involved in intercellular interactions between colonic and ileal infected immature enterocytes and resident immune cells. Bar plot showing the immune receptors targeted by upregulated (top graph) and downregulated (bottom graph) ligands in colonic (A) and ileal (B) infected immature enterocytes, scored by number of interactions (height of the bar plot) and number of immune cells targeted (black dots). The color of the bar plots indicates the number of ligands targeting each of the receptors indicated. This plot only shows the top 25 receptors by number of interactions, and the full plot is available as Supplementary Figure 6.

**Figure 6.**
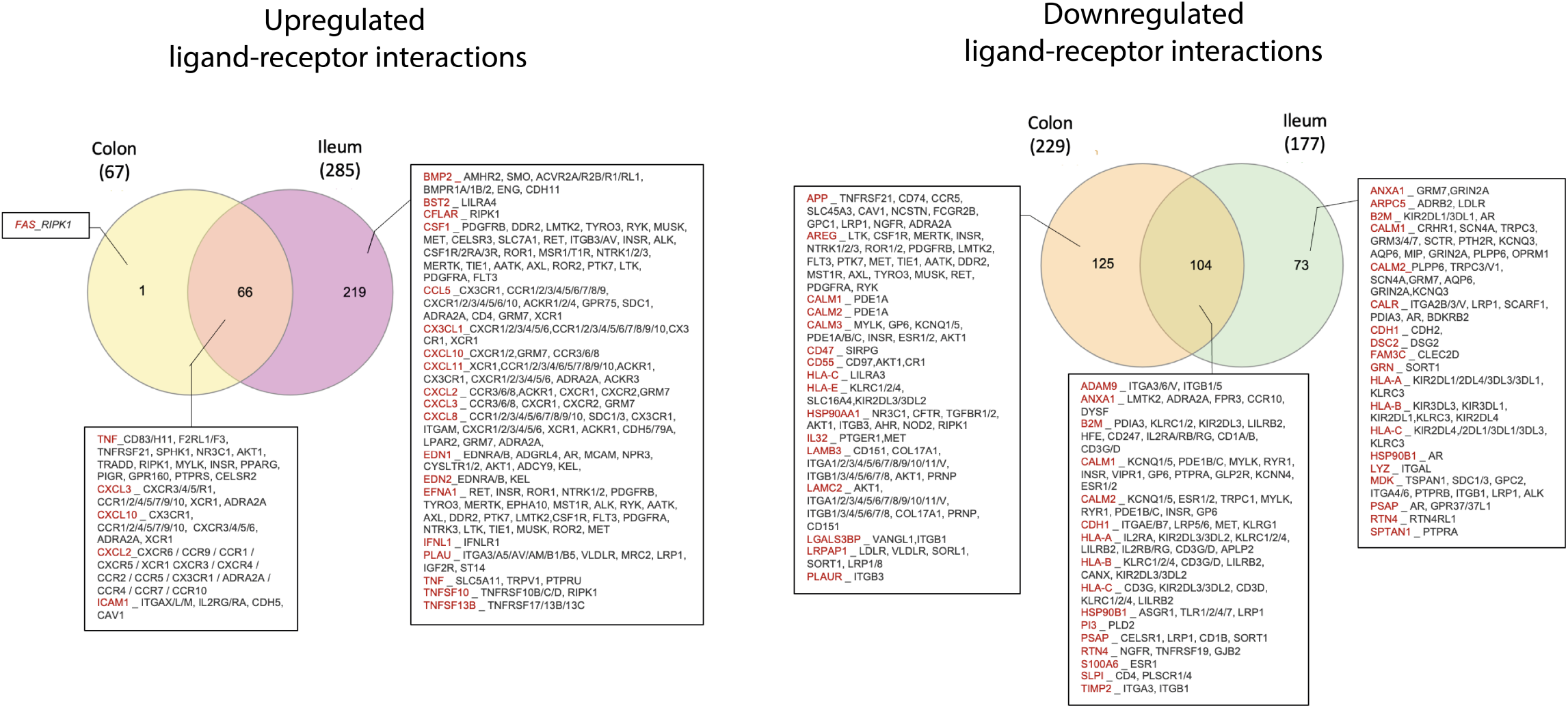
Overview of upregulated and downregulated ligand-receptor interactions between infected immature enterocytes and resident immune cells upon infection in the colon and ileum. Venn diagrams showing the number of ligand-receptor interactions in the infected immature enterocytes - immune cells intercellular network that are unique or shared between the ileum and colon. Intercellular interactions driven by upregulated and downregulated ligands are shown separately.

To understand which epithelial ligands and receptors were driving the strongest epithelial-immune cell interactions during infection, we scored ligands and receptors based on the “sum of receptor expression” value, which takes into account the number of interacting receptors and the level of receptor expression in each immune cell type (**Figure 1** **and Methods**). In the colon, the strongest upregulated interactions involved the epithelial TNF-a binding to B cells, T cells (CD4/CD8+), NK cells, macrophages and DCs, as well as epithelial chemokines (CXCL2,3, 10) binding to T cells (CD4/CD8+) and NK cells (**Figure 7A**). Similarly, in the ileum the strongest upregulated interactions involved epithelial chemokines binding to T cells (Treg, Tcyto, Tmem, CD8 Trm cyto) as well as TNF-a and CSF1 binding to macrophages and DCs (**Figure 7B**). Receptors driving the strongest upregulated interactions were mainly chemokine receptors (CXCRs, CCRs) in both colon and ileum (**Figure 9A, 9B**), and RIPK1 in the colon only (**Figure 9A**).

**Figure 7.**
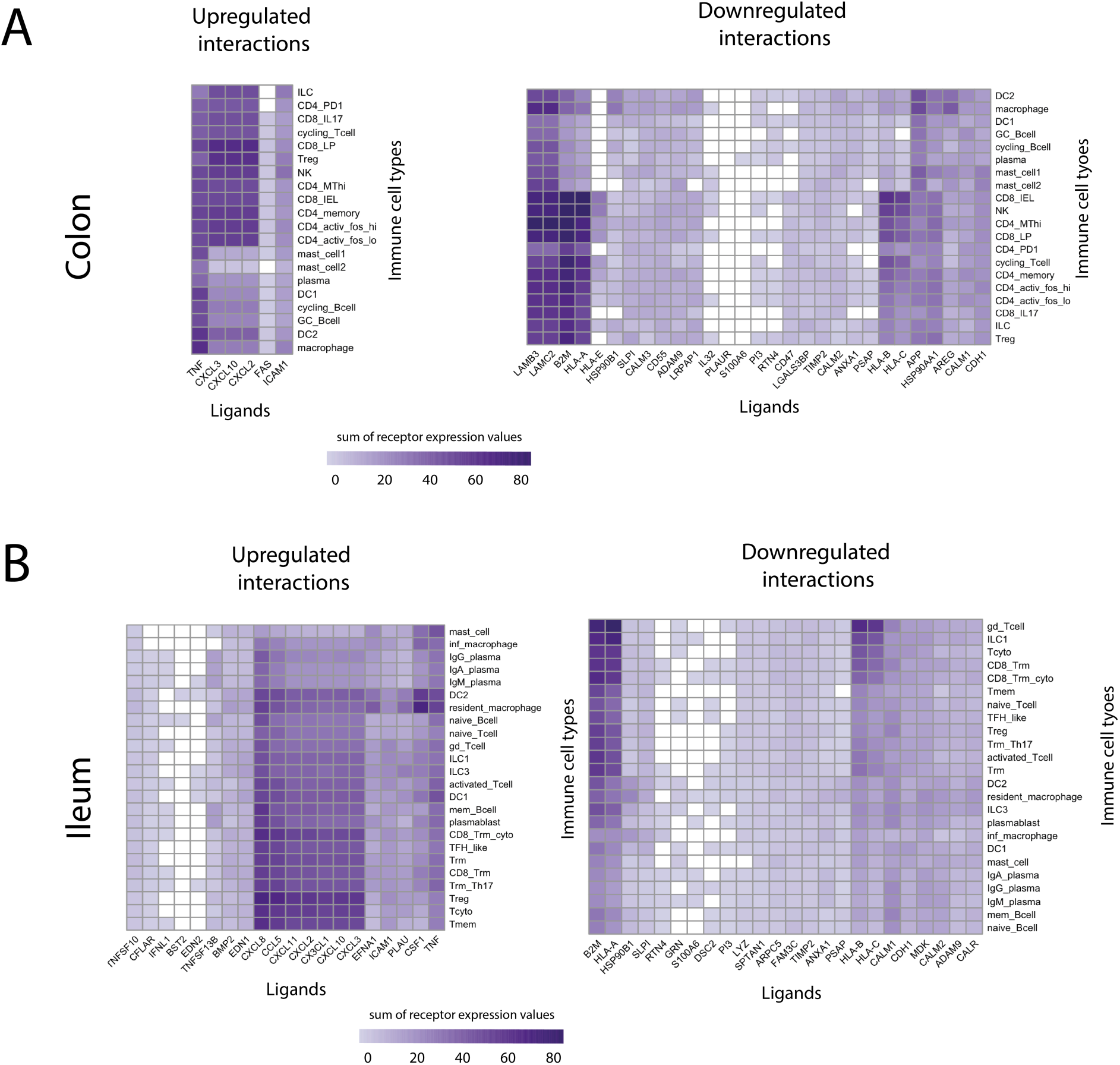
Ligands of infected immature enterocytes involved in the strongest up and downregulated interactions upon SARS-CoV-2 infection in the colon and ileum. **(A, B)** Heatplot showing the upregulated and downregulated interactions in the colon (A) and ileum (B) between intestinal epithelial ligands and resident immune cells upon infection of immature enterocytes with SARS-CoV-2. The strength of the interaction is expressed by accounting for the number of interactions between epithelial ligands and immune receptors and the level of receptor expression of immune cells. The strength of the interaction, named “sum of expression values”, is visualized using a color gradient from white (wekeast interactions) to purple (strongest interactions). Abbreviations: *<u>Ileum</u>*: inf_macrophage, infected macrophage; mast, mast cell, CD8_Trm_cyto, Resident memory cytotoxic T cell; DC2, dendritic cell 2; Trm, Tissue-resident memory T cell, gd_Tcell, Gamma delta (γδ) T cells; ILC, Innate lymphoid cell; mem_Bcell, memory B cell; naive_Bcell, naive B cell; TFH_like, T follicular helper cells; Trm_Th17, Tissue-resident memory Th17 cells; Treg, Regulatory T cell; Tcyto, Cytotoxic T cell; Tmem, Memory T cells. *<u>Colon</u>*: ILC, Innate lymphoid cell; CD8_IL17, IL-17+ CD8+ T cells; DC, dendritic cells; GC_Bcell, Germinal center B cells; CD4_PD1, mast, mast cell; Treg, Regulatory T cell; NK, Natural Killer cell, CD4_MThi, high mitochondrial CD4+ T cell; CD4_memory, CD4+ Memory T cell, CD4_activ_fos_high, activated CD4+ T cells (high/low c-fos); CD8_LP, CD8+ lymphocyte-predominant cells, CD8_IEL, CD8+ intraepithelial lymphocytes.

To understand the role of these epithelial-immune interactions, we performed a functional overrepresentation analysis of the participating upregulated epithelial ligands and receiving receptors on immune cells (**Figure 1, Supplementary Figure 8, and Methods**). In line with the extensive overlap in upregulated intercellular interactions (**Figure 6**), most functions were shared between colon and ileum, and included chemotaxis (GPCR signalling, chemokine signalling), immunity (interleukin signalling), apoptosis (caspase activation) and angiogenesis (VEGFA-VEGFR2 pathway) (**Supplementary Figure 9A, 9B**). One colonic-specific function was related to pro-inflammatory responses (TNF signalling) and one ileal-specific function was related to stem cell renewal (BMP signalling) **(Supplementary Figure 9A, 9B).**

### Downregulated epithelial ligands upon infection impact antigen presentation and focal adhesion pathways

Downregulated ligands in infected immature enterocytes were partially shared between colon and ileum, but were tissue-specific to a large extent (**Figure 2B**). Additionally, receptors on immune cells targeted by downregulated ligands were partially shared between colon and ileum (66), but several of them were tissue-specific (63 unique to colon, 38 unique to ileum) (**Figure 4**). In line with this, while some downregulated interactions in infected immature enterocytes were shared (104), a large proportion was tissue-specific (73 unique to ileum, 125 to colon) (**Figure 6** **& Supplementary Figure 8A, 8B).**

In both tissues, epithelial ligands involved in the highest number of interactions with immune cells included human leukocyte antigens (HLA-A/B/C)), beta-2-microglobulin (B2M) and calmodulin (CALM1/2), while receptors on immune cells included integrins (ITGs), Natural Killer Cell Lectin Like Receptors (KLRCs) and Killer Ig-like receptors (KIRs) (**Figure 2C, 2D, 6A, 6B**). Interestingly, the highest number of downregulated interactions in the colon upon infection was driven by two epithelial-derived laminins (LAMC2, LAMB3) and by integrins (ITGs) present on several different immune cell types (**Figure 2A, 6A**). Furthermore, receptors involved in the highest number of downregulated interactions were represented by AKT1 (Protein kinase B, PKB) uniquely in the colon, and by several integrins (ITGs), KLRCs and LDL Receptor Related Protein 1 (LRP1) in both colon and ileum (**Figure 5A, 5B**).

In the colon, ligands driving strong downregulated interactions were HLA-s (HLA-A, B, C) and B2M targeting CD4/CD8+ T cells, NK cells, ILCs and T reg cells (HLA-A and B2M only) (**Figure 7A**). In the ileum, we found similar downregulated interactions driven by HLAs and B2M, but the targeted immune cells included T cells (Trm, Tregs, cytotoxic T cell), ILCs (HLA-A, B, C only) and macrophages (both HLAs and B2M) (**Figure 7B**). Furthermore, the strongest downregulated interactions in the colon also included those among laminins (LAMB3, LAMC2) targeting T cells and macrophages, which we did not find in the ileum (**Figure 7A, 7B**). Receptors driving the strongest downregulated interactions were AKT1 uniquely in the colon (**Figure 8A**) as well as integrins, KLRCs and LRP1 in both colon and ileum (**Figure 8A, 8B**)

**Figure 8.**
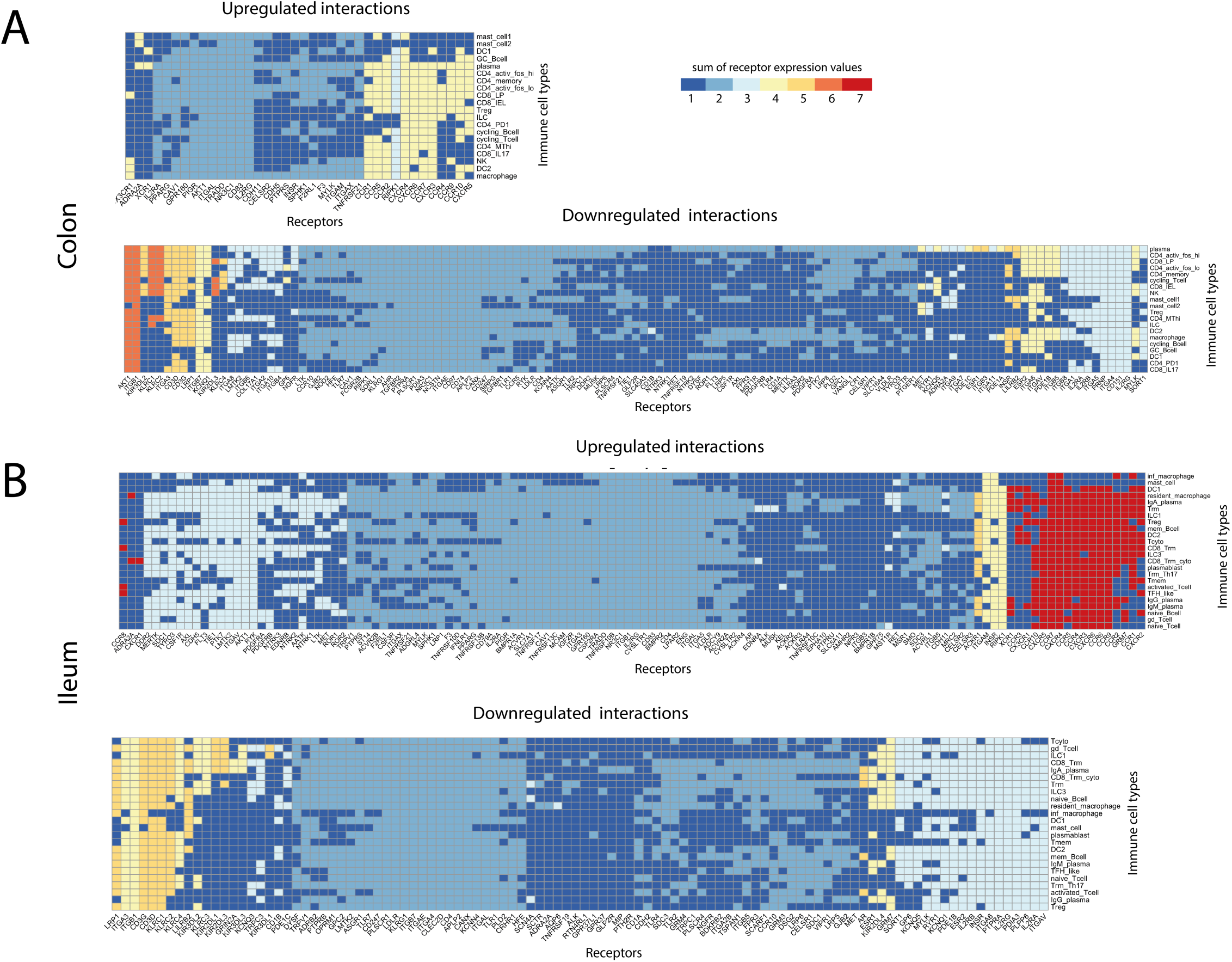
Receptors on immune cell types involved in the strongest up and downregulated interactions upon SARS-CoV-2 infection in the colon and ileum. **(A, B)** Heatplot showing the upregulated and downregulated interactions in the colon (A) and ileum (B) between receptors and resident immune cell types upon infection of immature enterocytes with SARS-CoV-2. The strength of the interaction is expressed by accounting for the number of interactions and the level of receptor expression of the receptor on immune cells. The strength of the interaction, named “sum of expression values”, is visualized using a color gradient from blue (wekeast interactions) to red (strongest interactions). Abbreviations: *<u>Ileum</u>*: inf_macrophage, infected macrophage; mast, mast cell, CD8_Trm_cyto, Resident memory cytotoxic T cell; DC2, dendritic cell 2; Trm, Tissue-resident memory T cell, gd_Tcell, Gamma delta (γδ) T cells; ILC, Innate lymphoid cell; mem_Bcell, memory B cell; naive_Bcell, naive B cell; TFH_like, T follicular helper cells; Trm_Th17, Tissue-resident memory Th17 cells; Treg, Regulatory T cell; Tcyto, Cytotoxic T cell; Tmem, Memory T cells. *<u>Colon</u>*: ILC, Innate lymphoid cell; CD8_IL17, IL-17+ CD8+ T cells; DC, dendritic cells; GC_Bcell, Germinal center B cells; CD4_PD1, mast, mast cell; Treg, Regulatory T cell; NK, Natural Killer cell, CD4_MThi, high mitochondrial CD4+ T cell; CD4_memory, CD4+ Memory T cell, CD4_activ_fos_high, activated CD4+ T cells (high/low c-fos); CD8_LP, CD8+ lymphocyte-predominant cells, CD8_IEL, CD8+ intraepithelial lymphocytes.

Functional overrepresentation analysis revealed shared functions related to antigen processing and cross-presentation (MHC class I–mediated), phagocytosis (ER phagosome pathway, Signalling by RHO GTPases) and cell-cell communication (immunoregulatory interactions between a lymphoid and non-lymphoid cell) in both tissues (**Supplementary Figure 9A, 9B**). Furthermore, several colon-specific functions were related to extracellular matrix organization and integrin cell surface interactions, suggesting decreased interactions involved in focal adhesion and intestinal tissue polarization uniquely in the colon (**Supplementary Figure 9A**). The only function uniquely overrepresented in the ileum was transcriptional regulation by MECP2 (**Supplementary Figure 9A**), whose expression has been shown to play a role in intestinal morphology and function (Millar-Büchner et al., 2016).

### Implication of epithelial ligands in the inflammatory process

With our experimental data-based analysis, we pointed out several differentially expressed ligands in the epithelial-immune cell network relative to infected immature enterocytes which could play a role in driving the inflammatory process upon SARS-CoV-2 infection. To validate their importance during immune reactions, we exploited independent data from three previously published studies (**Figure 1**).

First, by comparing the differentially expressed ligands upon SARS-CoV-2 infection to DEGs in human colonic organoids exposed to inflammatory cytokines (Pavlidis et al., 2021), we identified 24 ligands whose expression change is regulated by cytokines during intestinal inflammation (**Table 1, and Methods**). These ligands are more probable to contribute to the inflammatory responses upon infection. Next, by comparing ileal and colonic ligands to data from ImmunoGlobe, a manually curated intercellular immune interaction network (Atallah et al., 2020) and ImmunoeXpresso, a collection of cell–cytokine interactions generated through text mining (Kveler et al., 2018), we identified 12 ligands previously known to influence immune cell populations (**Table 1, and Methods**). The full list of affected immune cell types for each epithelial ligand is available in **Table 2B**. Finally, to understand which ileal and colonic ligands could explain blood cytokine level changes of COVID-19 patients via direct immune cell regulation, we used data from (Olbei et al., 2021), and identified 6 ligands capable to create the detected blood cytokine levels during infection (**Table 1, and Methods**).

**Table 1.** Key differentially expressed ligands produced by infected immature enterocytes drive the inflammatory process upon SARS-CoV-2 infection. Table showing a list of top-ranked differentially expressed ligands in infected immature enterocytes which were identified to drive inflammation upon SARS-CoV-2 infection. The ranking of the ligands was performed using multiple criteria as explained in the **Methods**. ‘Organoid type’ indicates whether the expression change of the ligand was found in ileal or colonic infected immature enterocytes upon SARS-CoV-2 infection, respectively. ‘Expression change upon SARS-CoV-2 infection’ indicates the direction of expression change of the ligand in infected immature enterocytes upon SARS-CoV-2 infection. ‘Regulation by cytokines’ indicates whether ligand expression was found to be regulated by cytokines during inflammation based on results from (Pavlidis et al., 2021). Ileal data was not available (n.d.) in this study, so no conclusions could be drawn for ileal ligands. ‘Known to affect immune cells’ indicates whether the ligand was found to be regulated by immune cells using data from ImmunoGlobe (1) and ImmunoeXpresso (2) databases. ‘Directly explain patient blood cytokine levels’ indicates whether the ligand was found to directly regulate blood cytokine levels in COVID-19 patients from (Olbei et al., 2021).

| Ranking | Ligand | Ligand description | Organoid type | Expression change upon SARS-CoV-2 infection | Regulation by cytokines | Known to affect immune cells | Directly explain patients blood cytokine levels |
| --- | --- | --- | --- | --- | --- | --- | --- |
| 1 | CXCL2 | C-X-C Motif Chemokine Ligand 2 | Colonic, Ileal | Up | IFNG, TNF, IL-17, IL-22 (up) | Neutrophils (1,2), fibroblasts, T cells, NK cell and CD8a+ DCs (1), leukocytes (2) | ✓ |
| 1 | CXCL3 | C-X-C Motif Chemokine Ligand 3 | Colonic, Ileal | Up | TNF, IL-22 (up) | Neutrophils, fibroblasts (1), T cells (2) | ✓ |
| 1 | CXCL10 | C-X-C Motif Chemokine Ligand 10 | Colonic, Ileal | Up | IFNG, TNF (up) | DC, Th1, NK cells, B cells, monocytes (1), and 29 additional immune cell types (2) | ✓ |
| 1 | CSF1 | Colony stimulating factor 1 | Colonic | Up |  | 35 immune cell types (2) | ✓ |
| 1 | CXCL11 | C-X-C Motif Chemokine Ligand 11 | Ileal | Up | n.d.* | DC (1,2), B cells, NK cells, Th1, monocytes, macrophages (1), lymphocytes, T cells, CD8+ alpha/beta T cell (2) | ✓ |
| 2 | TNFSF13B | TNF Superfamily Member 13b | Ileal | Up | n.d. | B cell, T cell, follicular B cell, naïve B cell, Th17, neutrophils, monocytes (2) | ✓ |
| 2 | LAMC2 | Laminin Subunit Gamma 2 | Colonic | Down | TNF, IL-22 (up) |  |  |
| 2 | CCL5 | C-C Motif Chemokine Ligand 5 | Ileal | Up | n.d.* | T cells, basophils, eosinophils, macrophages, monocytes, NK cells, DC, Memory T cells, Th1 and Th2 (1), and 23 additional immune cell types (2) |  |
| 2 | CX3CL1 | C-X3-C Motif Chemokine Ligand 1 | Ileal | Up | n.d.* | Monocytes, T cells, neutrophil, NK, DC, Mast cells and microglia (1), and 19 additional immune cell types (2) |  |
| 2 | CXCL8 | C-X-C Motif Chemokine Ligand 8 | Ileal | Up | n.d.* | Neutrophils, macrophages, basophils, naïve T cells, CD8+ T cells, monocytes (1), and 44 additional immune cell types (2) |  |
| 3 | ICAM1 | Intercellular Adhesion Molecule 1 | Colonic, Ileal | Up | IFNG, TNF, IL-22 (up) |  |  |
| 3 | IL32 | Interleukin 32 | Colonic | Down | IFNG, TNF (up) |  |  |
| 3 | AREG | Amphiregulin | Colonic | Down | IFNG (up), IL-13 (down) |  |  |
| 3 | TNF | Tumour Necrosis Factor | Colonic, Ileal | Up | TNF, IL-22 (+) | Non-specific: 129 immune cell types (2) |  |
| 3 | B2M | Beta-2-immunoglobulin | Colonic, Ileal | Down | IFNG (up) |  |  |
| 3 | HLA-A | Major Histocompatibility Complex, Class I, A | Colonic, Ileal | Down | IFNG (up) |  |  |
| 3 | HLA-B | Major histocompatibility complex, class I, B | Colonic, Ileal | Down | IFNG (up) |  |  |
| 3 | LAMB3 | Laminin Subunit Beta 3 | Colonic | Down |  |  |  |

\* Ileal data not available from Pavlidis *et al*, 2021
(1) Data from ImmunoGlobe (Atallah *et al*, 2020)
(2) Data from ImmunoeXpresso (Kveler *et al*, 2018)

Using this assessment, we were able to rank the differentially expressed ligands for their importance in the inflammatory process, and subsequently listed the 18 highest ranked ligands, for which there is strong evidence of their role in epithelial–immune cell interactions during the inflammatory SARS-CoV-2 disease response (**Table 1**). These ligands included CSF1, various chemokines (CXCL10, CXCL11, CXCL2, CXCL3, CCL5, CX3CL1, CXCL8), TNFa & TNFSF13B, and ICAM1 among the upregulated ones; and various laminins (LAMC2, LAMB3), AREG, B2M), human leukocyte antigens (HLAs) (HLA-A, HLA-B) and IL32 among the downregulated ones.

## Discussion

In this work, we have highlighted the putative central role of the gut during the immune response following SARS-CoV-2 infection, showing how several intracellular and intercellular mechanisms affected during infection, with key differences between colon and ileum.

SARS-CoV-2 has been shown to actively infect and reproduce in the human gut and in human gastro-intesinal tract derived organoids (Lamers et al., 2020; Stanifer et al., 2020; Triana et al., 2021a). However, it is not known what are the effects of intestinal inflammation and the role of epithelial-immune interactions in the hyperinflammatory immune response (“cytokine storm”) characterizing many COVID-19 patients (Arunachalam et al., 2020; Olbei et al., 2021). A better understanding of these interactions could help identify potential targets that are key to selectively disrupt such cell-cell interactions underlying extreme inflammatory conditions during SARS-CoV-2 infection. This would be extremely important given the failure of most randomized control trials associated with pro-inflammatory drug candidates for COVID-19 (Abubakar et al., 2020).

The altered epithelial-immune cell crosstalk during SARS-CoV-2 infection has been explored within the nasopharynx and lungs using scRNA seq data (Chua et al., 2020). This study found stronger epithelial-immune cell interactions in critically ill patients, based on ligand-receptor expression profiles, highlighting the importance of the crosstalk between infected cells and local immune cells in the disease course. However, to our knowledge no prior study has been carried out so far to model the effect of viral infection in host intestinal cells, and the role and contribution of intestinal epithelial cell–immune cell crosstalk during SARS-CoV-2 infection.

Our previous data (Triana et al., 2021a) has pointed out how immature enterocytes are the most affected epithelial cell population upon SARS-CoV-2 infection in the gut. Building on this knowledge, we developed a computational method to model and further investigate the effect of SARS-CoV-2 proteins and potential miRNAs on both ileal and colonic epithelial cells. We added in a distinguishable way the analysis of these potential miRNAs encoded by SARS-CoV-2 as previous studies highlighted the regulatory role of similar miRNAs produced by RNA viruses and their ability to downregulate host genes and affecting host functions (Bruscella et al., 2017; Griffiths-Jones et al., 2008; Saçar Demirci and Adan, 2020). In the present analysis, infected immature enterocytes were the most affected population based on the number of differentially expressed ligands, and drove most interactions with gut resident immune cells. Upon infection of immature enterocytes, several intracellular signalling pathways were altered, including those related to inflammation, apoptosis, cell survival and cell death (**Figure 3A, 3B**). Furthermore, pathways related to cell cycle (negative regulation of G2/M transition) and cell proliferation were also altered upon infection (**Figure 3A, 3B**), in line with a previous phosphoproteomics study finding a correlation with cell cycle arrest upon SARS-CoV-2 infection (Bouhaddou et al., 2020). Interestingly, several pathways involved in cell differentiation, cell migration and epithelial polarization, were also modulated upon infection in our study (**Figure 3A, 3B**).

By using available ligand-receptor interaction data, we aimed to elucidate mechanisms by which infected epithelial cells in the gut recruit innate and adaptive immune cell populations to find key interactions driving the immune response in the gut. In the colon, the number of downregulated ligands (29) was higher compared to upregulated ligands (6) upon infection (**Supplementary Figure 1**) and downregulated ligands drove the highest number of interactions (**Figure 2A**). Conversely, in the ileum, the number of upregulated (20) and downregulated ligands (24) was comparable (**Supplementary Figure 1)**, but upregulated ligands drove the highest number of cell-cell interactions with immune cells (**Figure 2A**). In both cases, the number of interactions with immune cells was not simply driven by the overall number of SARS-CoV-2 regulated ligands but by a few ligands presenting many different receptors on immune cells, and their relative expression change following infection (**Figure 2C, 2D & Supplementary Figure 8A, 8B**). For instance, in the colon, ligands driving the highest number of interactions included laminins (LAMB3, LAMC2) and MHC I - related proteins (HLA-A, HLA-B, B2M), which were all downregulated (**Figure 2A**). Conversely, in the ileum, the highest number of interactions was driven by cytokines/chemokines (TNF-a, CXCL8, CCL5, CXCL11) and adhesion factor CSF1, which are all upregulated (**Figure 2B**).

Most of upregulated ligands, mainly pro-inflammatory cytokines and chemokines, and cell-cell interactions they are involved in, are shared between colonic and ileal immature enterocytes upon infection, although more interactions are driven in the ileum (**Figure 2B** **&** **Figure 6**). Overall, upregulated interactions may reflect a general effect of the infection. Indeed, upregulated interactions were mainly represented by chemokines and TNF-a driven interactions (**Figure 2C, 2D & Figure 7A, 7B**), and functional analysis highlighted a relation to proinflammatory signalling pathways, including TNF-a signalling, interleukin signalling and chemotaxis *via* GPCR signalling, overall suggesting an increasing recruitment and cell adhesion of these immune cell populations upon infection (**Supplementary Figure 9A, 9B**). Notably, four chemokine receptors identified by our study (CXCR6 in the ileum, CCR1/2 and CCR9 in both ileum and colon) are coded in a genomic region that has been found associated as a COVID-19 risk locus on chromosome 3 (Schultze and Aschenbrenner, 2021). Importantly, recruitment of neutrophils by CXCL8 in the lung, which presented the epithelial ligand driving most interactions in the ileum in our study (**Figure 2D**), has been associated with disease severity in COVID-19 patients, supporting the role of chemokine-driven immune cell recruitment in disease manifestation (Park and Lee, 2020).

In both colon and ileum, we found strong downregulated interactions driven by epithelial HLAs (HLA-A, B, C) and B2M, a subcomponent of the major histocompatibility complex I (MHC I) (**Figure 2C, 2D**). According to our analysis, these ligands were mainly binding to KLR receptors, which are mainly presented on NK cells (**Supplementary Figure 8A, 8B**). Downregulation of HLAs represents a common immune evasion mechanism of viruses (Koutsakos et al., 2019), and has recently been discovered as a mechanism that SARS-CoV-2 protein ORF8 may use to escape host immune surveillance (Park, 2020).

Uniquely in the colon, we found strong downregulated interactions driven by epithelial laminins (LAMB3 and LAMC2) and integrins, with T cells and macrophages as the main immune cell types targeted upon infection (**Figure 2C, 6A**). Laminin-integrin binding contributes to focal adhesion of immune cells to the inflamed tissue (Simon and Bromberg, 2017) the overrepresentation of focal adhesion pathways and RHO GTPase signalling (**Supplementary Figure 9A**), which is involved in the migration of leukocytes to the site of infection (Biro et al., 2014). Overall, downregulation of laminins could represent a strategy for immune evasion following viral infection uniquely in the colon. Furthermore, laminins are known to play a role in shaping the architecture of intestinal mucosa, and an altered expression has been observed in Crohn’s disease, a type of IBD, driven by pro-inflammatory cytokines TNF-α and IFN-γ (Bouatrouss et al., 2000; Francoeur et al., 2004; Mahoney et al., 2008).

Finally, calmodulin genes (*CALM1, CALM2, CALM2*) were predicted to drive several downregulated ligand-receptor interactions (**Figure 2C, 3D**), mainly binding to cyclic AMP-specific phosphodiesterases (PDEs) (*PDE1A, PDE1B, PDE1C*) on immune cells in both tissues upon infection (**Supplementary Figure 8**). PDEs, whose activation is calcium/calmodulin dependent, are responsible for cyclic AMP (cAMP) degradation in T cells, which is a potent inhibitor of T-cell activation (Bjørgo et al., 2011). Hence, the downregulation of CALM-PDEs interactions following SARS-CoV-2 infection implies an increase in intracellular cAMP in T cells, and consequently an inhibition of their activity. This could represent another way during SARS-CoV-2 infection to evade the immune activation and viral clearance.

IgA plasma cells were the immune cell population with the highest number of cell-cell interactions upon infection in both colon and ileum (**Figure 2A**). Notably, previous reports suggests that IgA is the main type of immunoglobulin induced by mucosal infection of SARS-CoV-2, stressing the importance of the crucial role played by IgA-mediated mucosal immunity in anti-SARS-CoV-2 infection (Sterlin et al., 2021). Interestingly, in the colon most of these cell-cell interactions were driven by downregulated ligands, including laminins, HLAs and calmodulins, possibly suggesting a decreased antigen presentation and calcium-dependent activation of these cell types (**Figure 2A, 2C**). Conversely, in the ileum these interactions were driven by upregulated ligands, mainly cytokines/chemokines (TNF-a, CXCLs, CSF1) and adhesion factors (ICAM1, PLAU), possibly suggesting increased recruitment of these cell types to the epithelium (**Figure 2A, 2D**). The potential downregulation of cell-cell interactions with IgA plasma cells in the colon is an interesting avenue of further research. Of note, the size of each immune cell population was not taken into account in this analysis (**Methods**). Extension of the study in this way could refine the importance of each type of ligand-receptor communication in mediating the overall downstream functional changes, leading to a better prediction of the effect size for each ligand-receptor combination.

With our integrated workflow, we established a method to evaluate the effect of viral infection on host epithelial intestinal cells function and on the epithelial-immune crosstalk. Notably, this workflow is not limited to the gut, and it can be easily applied to other organs and cell types (e.g. lung, kidney, heart) which are relevant during SARS-CoV-2 infection and provided the right input data is available. Using our integrated intracellular and intercellular signalling network, we confirmed many of the previous findings about SARS-CoV-2 infection. These include the increase in pro-inflammatory responses, including TNF signalling and chemokine signalling, and the role played by T cells. Yet, we uncovered new mechanisms by which SARS-CoV-2 may evade the immune responses by interfering with epithelial-immune cell connections. Such mechanisms include downregulation of NK activation by HLAs-KLR interactions, focal adhesion pathways by laminin-integrins interactions, and T cell function by calmodulin-PDEs binding.

The methodology we used for our analysis has some limitations. When constructing the intracellular causal network, the effect of SARS-CoV-2 proteins towards human binding partners was always considered as inhibitory. However, this is not always the case. In the future, with increasingly available data, a more refined model could be generated. Furthermore, two different single cell transcriptomics datasets were used for colonic and ileal immune cell populations, due to the unavailability of both datasets from the same experiment. Similarly, IBD uninflamed data and healthy data were used for the ileum and colon respectively, as healthy control scRNAseq immune cell data for both tissues was not available at the time of the analysis. Finally, the *a priori* resources used to infer the intracellular and intercellular interaction networks may have some intrinsic limitations associated with them (Dimitrov et al., 2021) specific tools such as LIANA (LIgand-receptor ANalysis frAmework; https://github.com/saezlab/liana) could be used in the future to compare across several resources available, helping to choose the one(s) providing the best overall prediction.

With our analysis, we provided a set of intestinal epithelial ligands and immune cell populations implicated in altered epithelial-immune interactions during SARS-CoV-2 infection, which could potentially drive the excessive inflammatory processes seen in severe COVID-19 patients. Further experimental validation will be key to validate these processes and the key molecules and cell types involved. Introduction of immune cells to an organoid system is currently challenging. Yet, a recent study, where human intestinal CD4+ T cells have been co-cultured with human intestinal organoids (Schreurs et al., 2021), may represent a promising set-up for future studies to investigate epithelial-immune cell interactions during SARS-CoV-2 induced inflammation in the gut. As reviewed recently by (Min et al., 2020; Poletti et al., 2020), such co-culture systems could be excellent to study host-microbe interactions in the gut, including the detailed experimental analysis of SARS-CoV-2 infection in the gut.

With our work we presented a novel computational method to investigate the effect of SARS-CoV-2 proteins and miRNA on epithelial cell functions and epithelial-immune crosstalk upon infection. This workflow can be applied in the future to more epithelial and immune cell types when these data become available. Analyses of these intracellular and intercellular networks could shed light on the viral mechanisms of infection, the contribution of the gut to the cytokine storm, and possible location-dependent differences in effects of the viral infection.

## Acknowledgment

M.O., A.T., L.G., and M.P. are supported by the UKRI Biotechnological and Biosciences Research Council (BBSRC) funded Norwich Research Park Biosciences Doctoral Training Partnership (grant numbers BB/M011216/1 and BB/S50743X/1). The work of T.K. and D.M. was supported by the Earlham Institute (Norwich, UK) in partnership with the Quadram Institute (Norwich, UK) and strategically supported by the UKRI BBSRC UK grants (BB/J004529/1, BB/P016774/1, and BB/CSP17270/1). T.K. and D.M. were also funded by a BBSRC ISP grant for Gut Microbes and Health BB/R012490/1 and its constituent projects, BBS/E/F/000PR10353 and BBS/E/F/000PR10355. T.A. and S.T. acknowledge the funding from the Darwin Trust of Edinburgh and from the ERC Consolidator grant METACELL from European Union’s Horizon 2020 program. T.A. and S.T. acknowledge support from the EMBL Genomics Core Facility and particularly help from Vladimir Benes. B.V. is supported by the Clinical Research Fund (KOOR) University Hospitals Leuven. S.B. was supported by research grants from the Deutsche Forschungsgemeinschaft (DFG): project numbers 415089553 (Heisenberg program), 240245660 (SFB1129), 278001972 (TRR186), and 272983813 (TRR179), the state of Baden Wuerttemberg (AZ: 33.7533.-6-21/5/1) and the Bundesministerium Bildung und Forschung (BMBF) (01KI20198A). M.L.S. was supported by the DFG (416072091) and the BMBF (01KI20239B). D.T. was supported by the Federal Ministry of Education and Research (BMBF, Computational Life Sciences grant no. 031L0181B) to J.S.R.

## Supplementary Figures

**Supplementary Figure 1. Differentially expressed ligands upon SARS-CoV-2 infection in infected or bystander epithelial sub-populations**. Bar chart indicating the number of differentially expressed ligands in the intercellular network in each epithelial sub-populations, either bystander or infected, in ileal or colonic organoids infected with SARS-CoV-2 vs control (24 hrs). Differentially expressed ligands are those DEGs found in (Lamers et al., 2020; Stanifer et al., 2020; Triana et al., 2021a; Zang et al., 2020), for which at least one binding receptor was found on immune cell populations. Color of the bar indicates the direction of regulation (red, upregulated; blue: downregulated). Abbreviations: ta, transit amplifying; imm_enterocyte, immature enterocyte.

**Supplementary Figure 2. Intracellular signalling networks of ileal and colonic infected immature enterocytes upon SARS-CoV-2 infection.** A, B) Causal networks of SARS-CoV-2-infected colonic and ileal immature enterocytes reconstructed using ViralLink or CARNIVAL. For each network the number of interacting partners (nodes), number of interactions (edges), average number of neighbours, network diameter and characteristics path length are indicated under “Network characteristics”. Within the “Node table”, columns indicate the types of nodes in the different layers of the network, including SARS-CoV-2 proteins or miRNAs, human binding proteins, intermediary signalling proteins, transcription factors (TFs) and differentially expressed ligands. Where an interacting partner (human protein/gene) was found to act in multiple layers of the network, it was assigned to a layer based on the following priority: differentially expressed ligands, human binding proteins, TFs, intermediary signalling proteins. Ligands have log2 fold change ≥ |0.5| and adjusted p value ≤ 0.05. Within the “Node Table”, rows indicate whether the node or edge belong to intracellular signals stemming from viral miRNA only, viral protein only or both (“shared”).

**Supplementary Figure 3. Overview of intracellular signalling upon SARS-CoV-2 infection in colonic infected immature enterocytes, reconstructed using the CARNIVAL**. From left to right: signalling cascade going from the upstream perturbation (SARS-CoV-2 proteins or miRNAs interacting with human binding proteins) to the downstream perturbation, transcription factors (TFs) regulating the differentially expressed ligands. Diamonds indicate the most active transcription factors after infection and the ovals are the perturbed human binding proteins. Rectangles are signaling intermediate proteins linking these two. Parallelograms and downward arrows indicate SARS-CoV-2 proteins and miRNAs, respectively. The color of the node indicates activation (red) or inhibition (blue) upon SARS-CoV-2 infection vs uninfected condition. Connecting edges show the direction of the interaction, as activation (pointed arrow) or inhibition (T shape arrow). Differentially expressed ligands for which no upstream signalling was identified, but downstream intercellular connections were predicted are excluded from this figure. Differentially expressed ligands are grouped based on the direction of regulation, which is indicated with blue when downregulated (bottom) and red when upregulated (top) when comparing SARS-CoV-2 infected vs uninfected conditions. Colors of the nodes edge and of the functional analysis boxes indicate if the original network was a miRNA only (yellow), viral protein only (black) or both viral protein and miRNA (grey). Functional overrepresentation analysis was carried out for the “PPI layer” of the intracellular network which includes human binding proteins, intermediary signalling proteins and TFs (adj p value < 0.05, n > 3).

**Supplementary Figure 4. Overview of intracellular signalling upon SARS-CoV-2 infection in ileal infected immature enterocytes, reconstructed using CARNIVAL.** From left to right: signalling cascade going from the upstream perturbation (SARS-CoV-2 proteins or miRNAs interacting with human binding proteins) to the downstream perturbation, transcription factors (TFs) regulating the differentially expressed ligands. Diamonds indicate the most active transcription factors after infection and the ovals are the perturbed human binding proteins. Rectangles are signaling intermediate proteins linking these two. Parallelograms and downward arrows indicate SARS-CoV-2 proteins and miRNAs, respectively. The color of the node indicates activation (red) or inhibition (blue) upon SARS-CoV-2 infection vs uninfected condition. Connecting edges show the direction of the interaction, as activation (pointed arrow) or inhibition (T shape arrow). Differentially expressed ligands for which no upstream signalling was identified, but downstream intercellular connections were predicted are excluded from this figure. Differentially expressed ligands are grouped based on the direction of regulation, which is indicated with blue when downregulated (bottom) and red when upregulated (top) when comparing SARS-CoV-2 infected vs uninfected conditions. Colors of the nodes edge and of the functional analysis boxes indicate if the original network was a miRNA only (yellow), viral protein only (black) or both viral protein and miRNA (grey). Functional overrepresentation analysis was carried out for the “PPI layer” of the intracellular network which includes human binding proteins, intermediary signalling proteins and TFs (adj p value < 0.05, n > 3).

**Supplementary Figure 5. Differentially expressed ligands of colonic and ileal bystander immature enterocytes upon SARS-CoV-2 infection.** Bar plot showing the upregulated and downregulated ligands in the colonic (top) and ileal (bottom) bystander immature enterocytes - immune cell network scored by number of interactions (height of the bar plot) and number of immune cells targeted (black dots). Upregulated ligands are shown in red and downregulated ligands in blue.

**Supplementary Figure 6. Receptors involved in intercellular interactions between colonic and ileal infected immature enterocytes and resident immune cells.** Bar plot showing the immune receptors targeted by upregulated (top graph) and downregulated (bottom graph) ligands in colonic (A) and ileal (B) infected immature enterocytes, scored by number of interactions (height of the bar plot) and number of immune cells targeted (black dots). The color of the bar plots indicates the number of ligands targeting each of the receptors indicated.

**Supplementary Figure 7. Receptors involved in intercellular interactions between colonic and ileal bystander immature enterocytes and resident immune cells.** Bar plot showing the immune receptors targeted by upregulated (top graph) and downregulated (bottom graph) ligands in colonic (A) and ileal (B) bystander immature enterocytes, scored by number of interactions (height of the bar plot) and number of immune cells targeted (black dots). The color of the bar plots indicates the number of ligands targeting each of the receptors indicated.

**Supplementary Figure 8. Intercellular interactions with upregulated and downregulated ligands of colonic and ileal infected immature enterocytes**. Interactions driven by upregulated and downregulated ligands are shown separately. The number of immune cells involved in ligand-receptor interaction pair is indicated in purple.

**Supplementary Figure 9. Functional analysis of ligand-receptor interactions between ileal and colonic immature enterocytes and resident immune cells upon SARS-CoV-2. A, B)** Reactome functional overrepresentation analysis carried out a list of all upregulated ligands and receptors for interactions of a specific condition. There was no weighting for the number of interactions of each ligand/receptor. Analyses relative to interactions driven by upregulated and downregulated ligands are shown separately.

**Supplementary Figure 10. Functional analysis of ligand-receptor interactions between ileal and colonic immature enterocytes and resident immune cells upon SARS-CoV-2.** Reactome functional overrepresentation analysis carried out a list of all upregulated ligands and receptors for interactions of a specific condition. There was no weighting for the number of interactions of each ligand/receptor. For the colon, analyses relative to interactions driven by upregulated and downregulated ligands are shown separately. For the ileum, analyses relative to upregulated ligands only are shown, as there were no interactions driven by downregulated ligands.

