## Supplementary Text for "Reprogramming of the intestinal epithelial-immune cell interactome during SARS-CoV-2 infection"

### Supplementary Information

Intracellular signalling network upon SARS-CoV-2 infection

The colonic network was made of 1423 nodes and 9971 edges, and had a network diameter of 10, characteristic path length of 4 and average number of neighbours of 14 (**Table 1**). Additionally, the ileal network was made of 1316 nodes and 7935 edges, and had a network diameter of 9, characteristic path length of 4 and average number of neighbours of 12 (**Table 1**). In the colon, we found 47 viral proteins/miRNAs, 409 human binding proteins, 908 intermediary signalling proteins, 37 TFs and 22 differentially expressed ligands, while in the ileum we found 47 viral proteins or miRNAs, 394 human binding proteins, 810 intermediary signalling proteins, 37 TFs and 28 differentially expressed ligands **(Table 1).**

In the colon, following SARS-CoV-2 infection, pathways related to chemotaxis, cell motility and extracellular/laminin interactions were overrepresented uniquely in the viral miRNA network. Additionally, we found functions related to inflammation (NF-kB, IL-1), cell cycle (negative regulation of G2/M transition) and establishment of tissue polarity (WNT signalling, Hedgehog signalling) overrepresented uniquely in the viral protein network (**Figure 3A**). Finally, we found functions related to cell growth (ESR, VEGF), negative regulation of apoptosis, positive regulation of cell migration, proliferation and differentiation (including SMAD/TGF-b signalling), and inflammation (cytokines, interleukins signalling) overrepresented in both viral miRNA and protein networks (**Figure 3A**). Similar results were found in the ileum, with the addition of functions involved in T cell activation and interferon gamma (IFN-γ) signalling overrepresented uniquely in the viral miRNA network and MAPK signalling overrepresented in both viral miRNA and protein networks **(Figure 3B).**

To better understand the main mechanisms responsible for the changes in the expression levels observed in infected immature enterocytes, we looked at transcriptional regulators directly regulating the differentially expressed ligands in immature enterocytes of both colon and ileum. In the colon, differentially expressed ligands were regulated upstream by TFs with a role in cell survival, proliferation and death (ATF2/3, ETS1/2, EGR1, JUN/JUNB, JUND (AP-1), FOS), interferon signalling (IRF1, STAT1), hypoxia (HIF1A) and intestinal cell differentiation (KLF4), which were all upregulated (not shown, full network uploaded on the GitHub repository of the project). In the ileum, we found very similar results, with the addition of FOXO (FOXO1/3/4) and SMAD3, both involved in intestinal cell differentiation, which were also upregulated (not shown, full network uploaded on the GitHub repository of the project). When using a different method (CARNIVAL) to build the intracellular network, we could find very comparable results (**Supplementary Figure 3,4**).

Epithelial-immune interactome driven by SARS-CoV-2 regulated ligands bystander epithelial cells upon infection

In bystander cells, the number of ligands was much lower than in infected cells (**Supplementary Figure 1**). Nevertheless, similar effects could be found, with the highest number of upregulated interactions driven by epithelial chemokines (CXCL10/11) and TNF-a binding to chemokine receptors (CXCRs, CCRs) and TNF receptors (ileum only) on immune cells in both tissues (**Supplementary Figure 5A, 5B, 7A, 7B)**. Strong upregulated interactions involved epithelial chemokines and various CD4+ and CD8+ T cells, B cells and ILCs in both colon and ileum **(Supplementary Figure 11A, 11B).** Functional overrepresentation analysis revealed that these interactions were mainly related to recruitment of immune cells to the epithelium (GPCR signalling, chemokine signalling) and cell death/necrosis in both colon and ileum (**Supplementary Figure 10A, 10B**).

In bystander cells, most downregulated interactions were driven by epithelial laminin (LAMA3) binding to integrins (ITGs) on immune cells in the colon only (**Supplementary Figure 5A, 7A**), while no downregulated interactions were found in the ileum (**Supplementary Figure 5B, 7B**). In the colon, strongest downregulated interactions involved epithelial laminin LAMA3 and various CD4+ and CD8+ T cells, NK cells and macrophages **(Supplementary Figure 11A**). Functional overrepresentation analysis revealed that these interactions were mainly related to extracellular matrix organization (**Supplementary Figure 10A).**
