## Supplementary Figures (S1-11) for "Reprogramming of the intestinal epithelial-immune cell interactome during SARS-CoV-2 infection"

### Figure S1

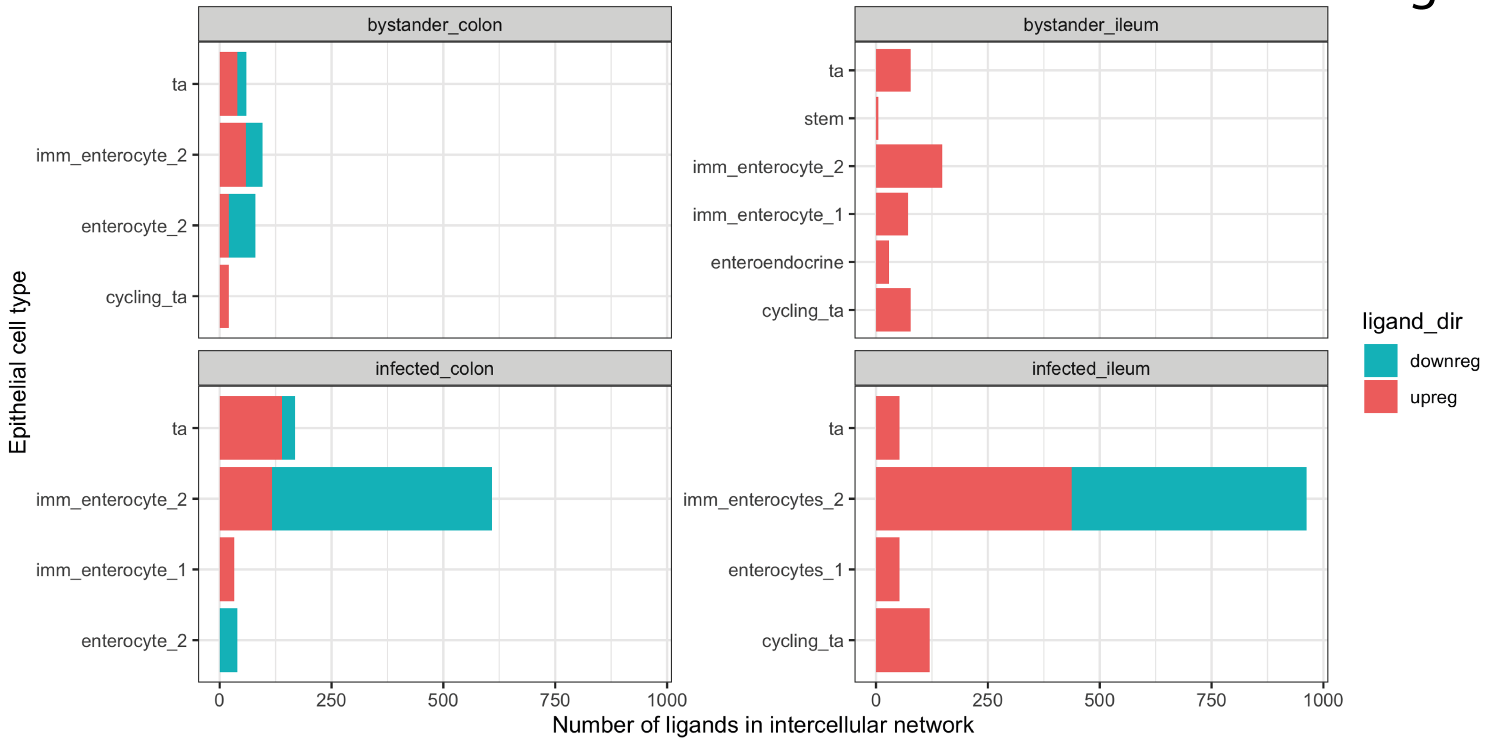

bystander_colon
bystander_ileum
infected_colon
infected_ileum
ta
imm_enterocyte_2
enterocyte_2
cycling_ta
stem
imm_enterocyte_1
enteroendocrine
imm_enterocytes_2
enterocytes_1
0
250
500
750
1000
Epithelial cell type
Number of ligands in intercellular network
ligand_dir
downreg
upreg

### Figure S2

#### ViralLink

##### Colon

###### Network characteristics

| Network parameter | Statistics |
|---|---|
| Number of nodes | 1423 |
| Number of edges | 9971 |
| Average number of neighbours | 14 |
| Network diameter | 10 |
| Path length | 4 |

###### Node table

| | Viral miRNAs or proteins | Human Binding proteins | Intermediary signalling proteins | TFs | Differentially expressed ligands | **Total** |
|---|---|---|---|---|---|---|
| Viral miRNA specific | 19 | 107 | 120 | 4 | 2 | **252** |
| Viral protein specific | 28 | 236 | 223 | 4 | 2 | **493** |
| Shared | 0 | 66 | 565 | 29 | 18 | **678** |
| **Total** | **47** | **409** | **908** | **37** | **22** | **1423** |

##### Ileum

###### Network characteristics

| Network parameter | Statistics |
|---|---|
| Number of nodes | 1316 |
| Number of edges | 7935 |
| Average number of neighbours | 12 |
| Network diameter | 9 |
| Path length | 4 |

###### Node table

| | Viral miRNAs or proteins | Human Binding proteins | Intermediary signalling proteins | TFs | Differentially expressed ligands | **Total** |
|---|---|---|---|---|---|---|
| Viral miRNA specific | 28 | 125 | 242 | 6 | 1 | **402** |
| Viral protein specific | 19 | 214 | 169 | 2 | 1 | **405** |
| Shared | 0 | 54 | 399 | 29 | 26 | **508** |
| **Total** | **47** | **393** | **810** | **37** | **28** | **1315** |

#### CARNIVAL

##### Colon

###### Network characteristics

| Network parameter | Statistics |
|---|---|
| Number of nodes | 147 |
| Number of edges | 279 |
| Average number of neighbours | 3.782 |
| Network diameter | 15 |
| Path length | 6.375 |

###### Node table

| | Viral miRNAs or proteins | Human Binding proteins | Intermediary signalling proteins | TFs | Differentially expressed ligands | **Total** |
|---|---|---|---|---|---|---|
| Viral miRNA specific | 4 | 0 | 31 | 22 | 0 | **57** |
| Viral protein specific | 16 | 25 | 4 | 10 | 0 | **55** |
| Shared | 0 | 2 | 1 | 11 | 21 | **35** |
| **Total** | **20** | **27** | **36** | **43** | **21** | **147** |

##### Ileum

###### Network characteristics

| Network parameter | Statistics |
|---|---|
| Number of nodes | 274 |
| Number of edges | 479 |
| Average number of neighbours | 3.219 |
| Network diameter | 9 |
| Path length | 3.353 |

###### Node table

| | Viral miRNAs or proteins | Human Binding proteins | Intermediary signalling proteins | TFs | Differentially expressed ligands | **Total** |
|---|---|---|---|---|---|---|
| Viral miRNA specific | 4 | 0 | 154 | 1 | 3 | **162** |
| Viral protein specific | 17 | 30 | 15 | 0 | 1 | **61** |
| Shared | 0 | 1 | 0 | 25 | 21 | **49** |
| **Total** | **21** | **31** | **169** | **26** | **25** | **272** |

### Figure S3

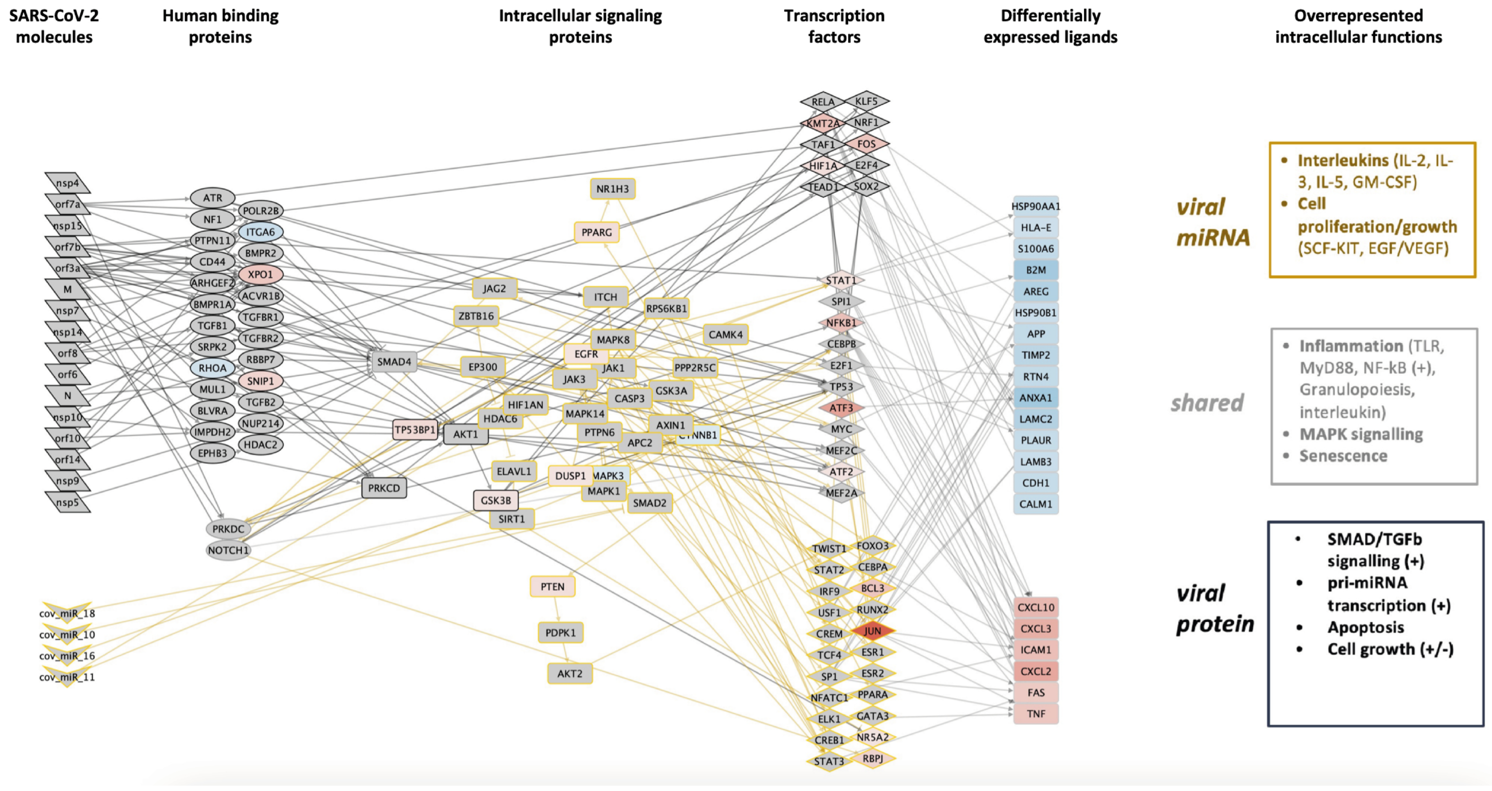


Colon

### Figure S4

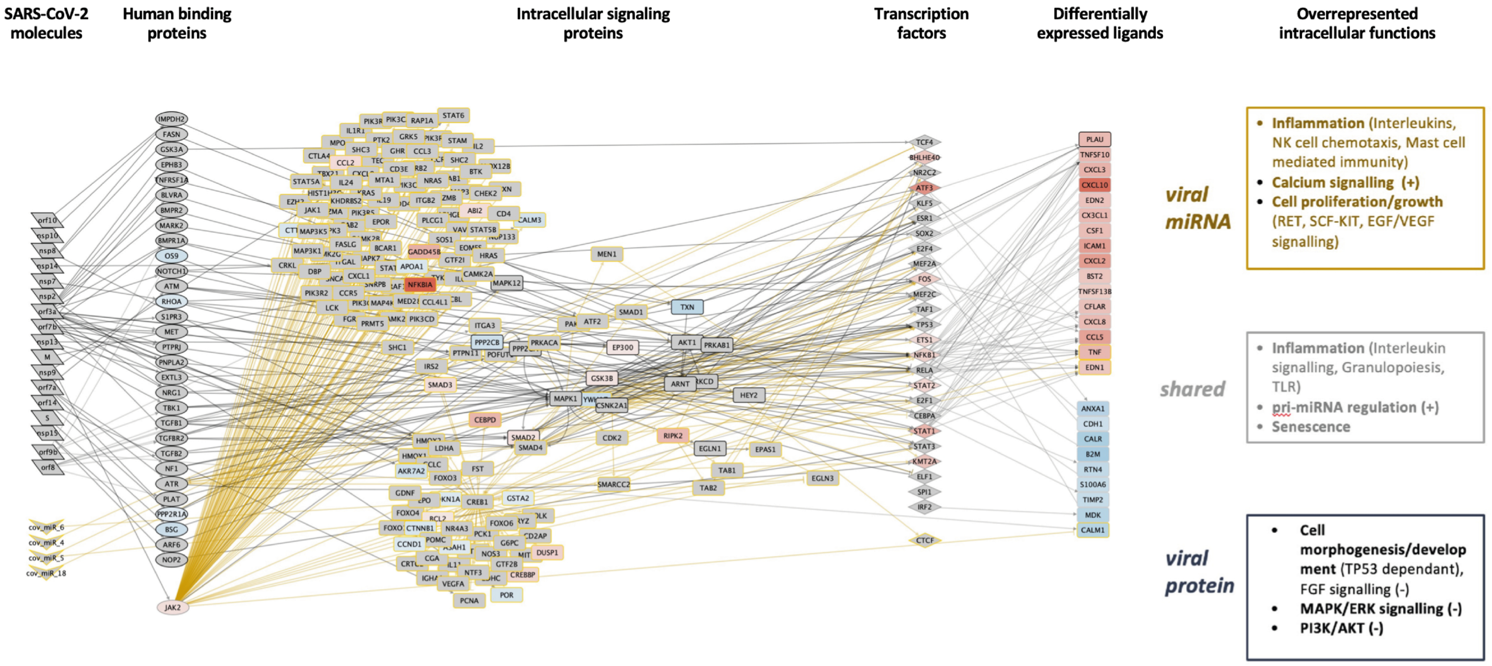


Ileum

### Figure S5

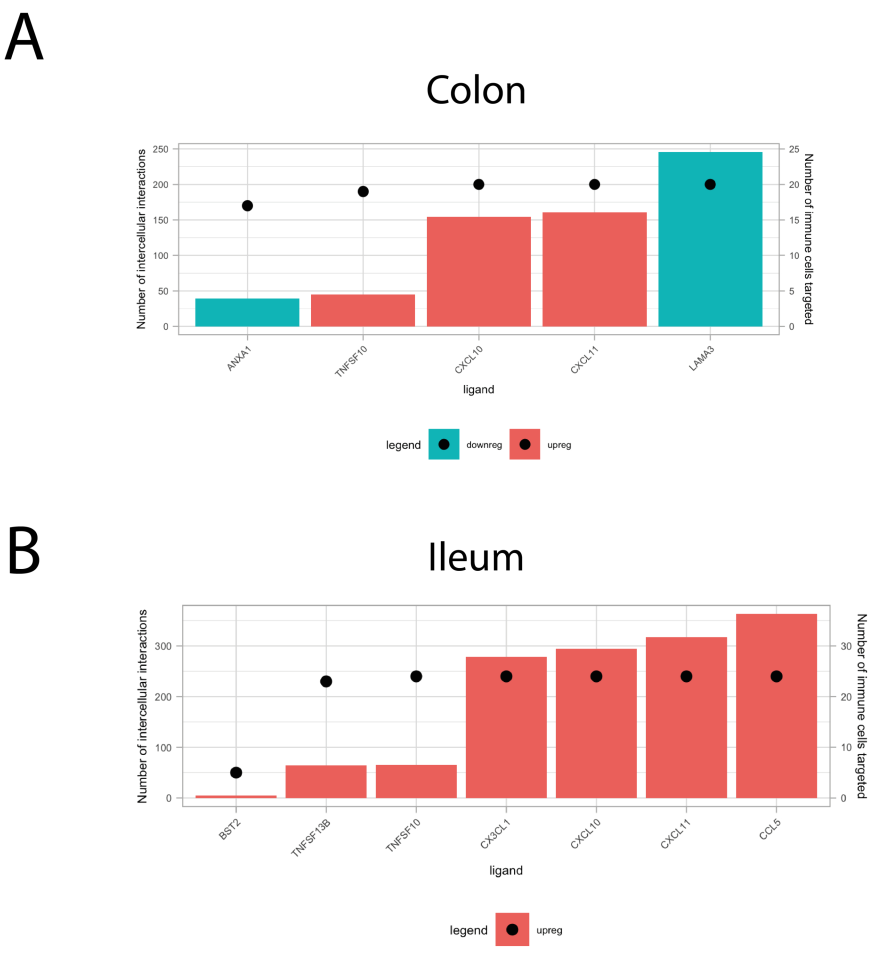
A
Colon
Number of intercellular interactions
Number of immune cells targeted
ANXA1
TNFSF10
CXCL10
CXCL11
LAMA3
ligand
legend
downreg
upreg
B
Ileum
Number of intercellular interactions
Number of immune cells targeted
BST2
TNFSF13B
TNFSF10
CX3CL1
CXCL10
CXCL11
CCL5
ligand
legend
upreg

Figure S6

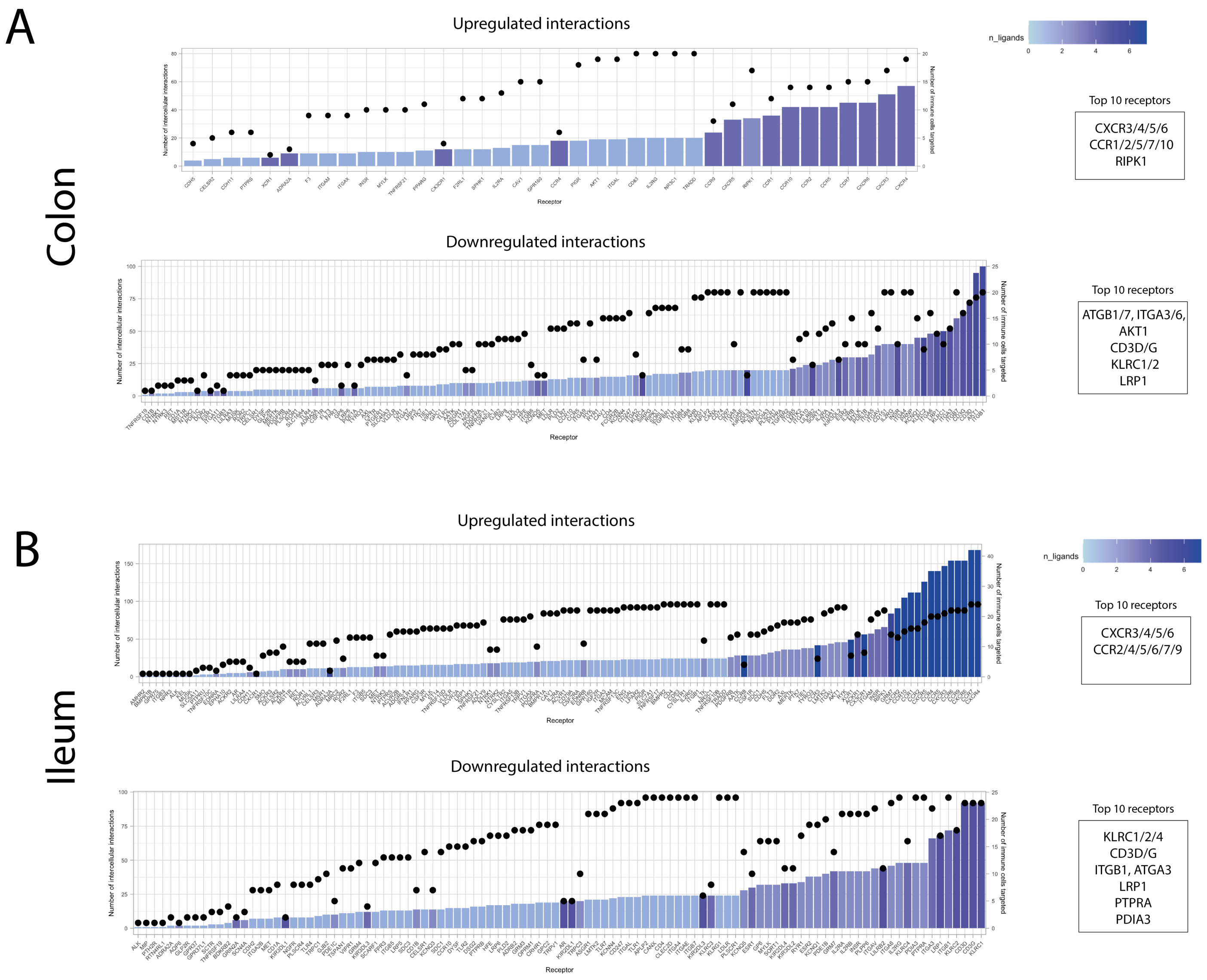
A
Colon
Upregulated interactions
Number of intercellular interactions
Number of immune cells targeted
Receptor
n_ligands
Top 10 receptors
CXCR3/4/5/6
CCR1/2/5/7/10
RIPK1
Downregulated interactions
Number of intercellular interactions
Number of immune cells targeted
Receptor
Top 10 receptors
ATGB1/7, ITGA3/6,
AKT1
CD3D/G
KLRC1/2
LRP1
B
Ileum
Upregulated interactions
Number of intercellular interactions
Number of immune cells targeted
Receptor
n_ligands
Top 10 receptors
CXCR3/4/5/6
CCR2/4/5/6/7/9
Downregulated interactions
Number of intercellular interactions
Number of immune cells targeted
Receptor
Top 10 receptors
KLRC1/2/4
CD3D/G
ITGB1, ATGA3
LRP1
PTPRA
PDIA3

### Figure S7

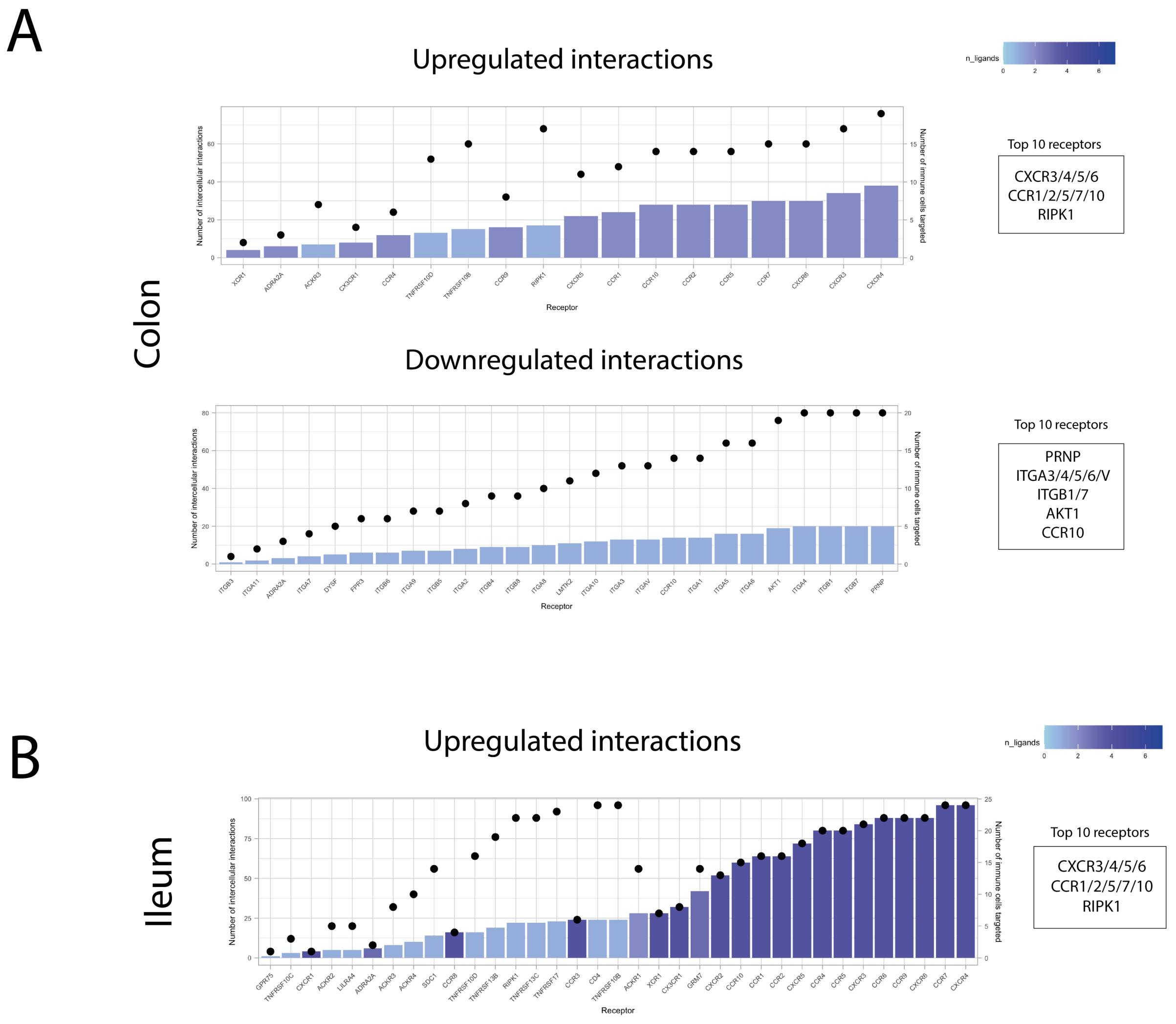


Bystander immature enterocytes

#### Figure S8

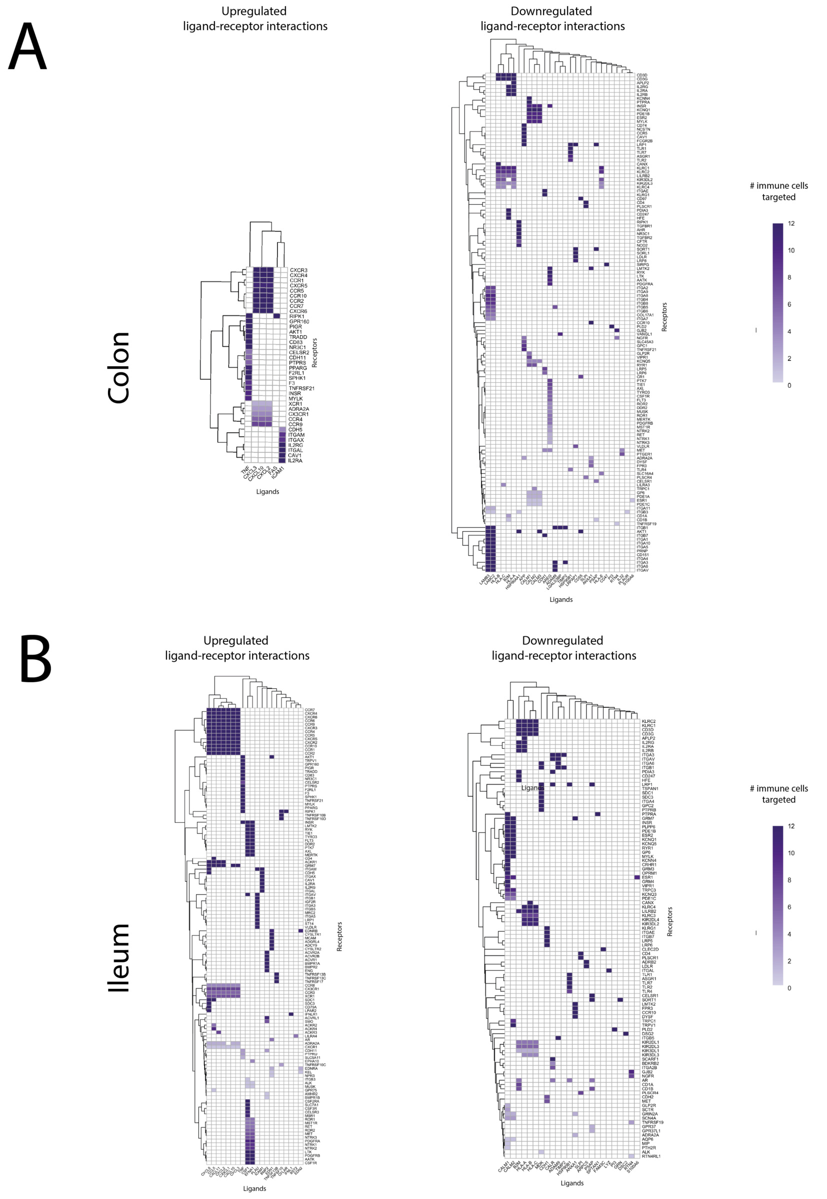

A
Upregulated
ligand-receptor interactions
Downregulated
ligand-receptor interactions
Colon
Ligands
Receptors
### immune cells
targeted
12
10
8
6
4
2
0
B
Upregulated
ligand-receptor interactions
Downregulated
ligand-receptor interactions
Ileum
Ligands
Receptors
### immune cells
targeted
12
10
8
6
4
2
0

### Figure S9

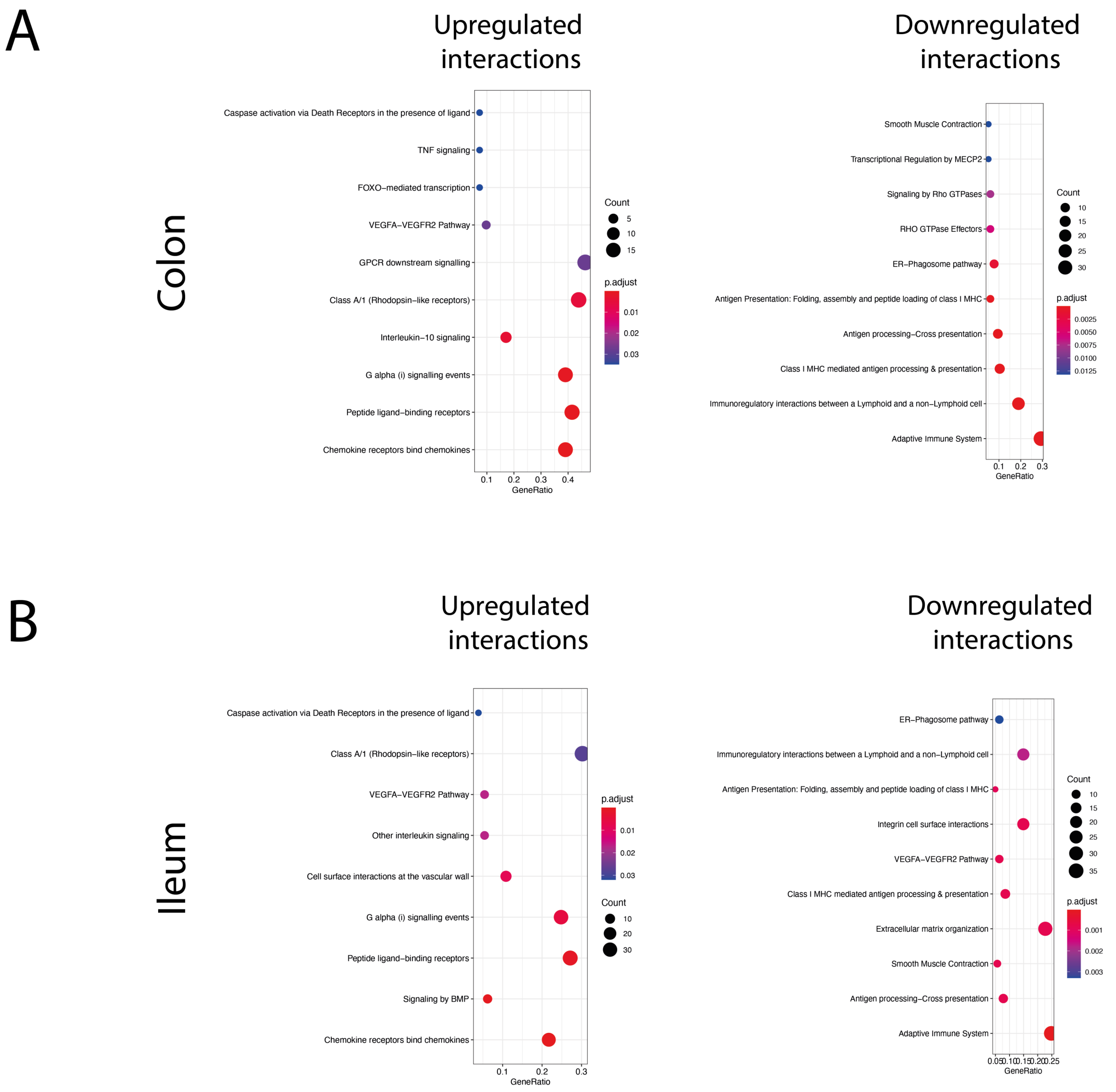

A
Colon
Upregulated interactions
Caspase activation via Death Receptors in the presence of ligand
TNF signaling
FOXO–mediated transcription
VEGFA–VEGFR2 Pathway
GPCR downstream signalling
Class A/1 (Rhodopsin–like receptors)
Interleukin–10 signaling
G alpha (i) signalling events
Peptide ligand–binding receptors
Chemokine receptors bind chemokines
GeneRatio
Count
p.adjust
Downregulated interactions
Smooth Muscle Contraction
Transcriptional Regulation by MECP2
Signaling by Rho GTPases
RHO GTPase Effectors
ER–Phagosome pathway
Antigen Presentation: Folding, assembly and peptide loading of class I MHC
Antigen processing–Cross presentation
Class I MHC mediated antigen processing & presentation
Immunoregulatory interactions between a Lymphoid and a non–Lymphoid cell
Adaptive Immune System
GeneRatio
Count
p.adjust
B
Ileum
Upregulated interactions
Caspase activation via Death Receptors in the presence of ligand
Class A/1 (Rhodopsin–like receptors)
VEGFA–VEGFR2 Pathway
Other interleukin signaling
Cell surface interactions at the vascular wall
G alpha (i) signalling events
Peptide ligand–binding receptors
Signaling by BMP
Chemokine receptors bind chemokines
GeneRatio
p.adjust
Count
Downregulated interactions
ER–Phagosome pathway
Immunoregulatory interactions between a Lymphoid and a non–Lymphoid cell
Antigen Presentation: Folding, assembly and peptide loading of class I MHC
Integrin cell surface interactions
VEGFA–VEGFR2 Pathway
Class I MHC mediated antigen processing & presentation
Extracellular matrix organization
Smooth Muscle Contraction
Antigen processing–Cross presentation
Adaptive Immune System
GeneRatio
Count
p.adjust

### Figure S10

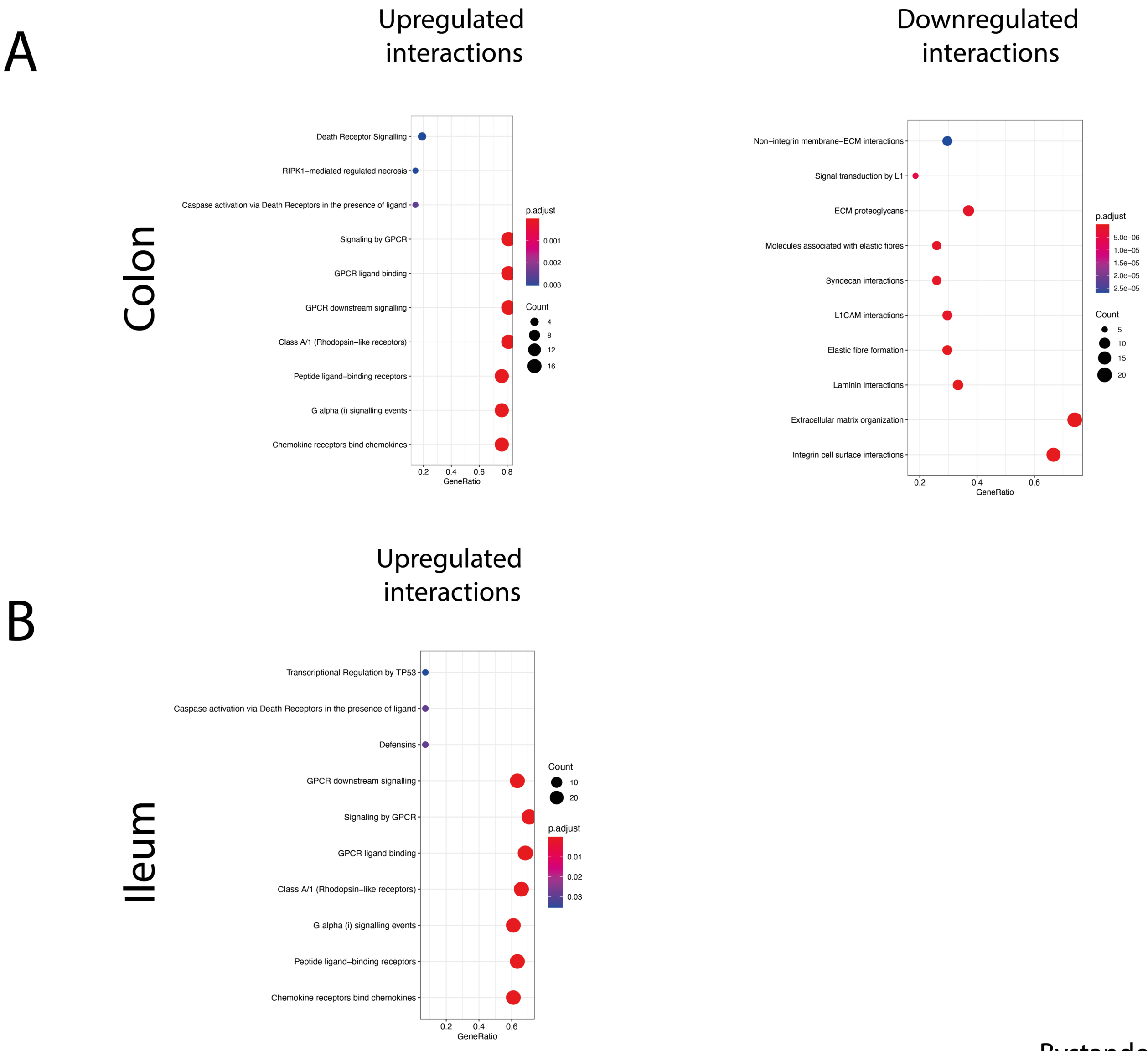


Bystander immature enterocytes

### Figure S11

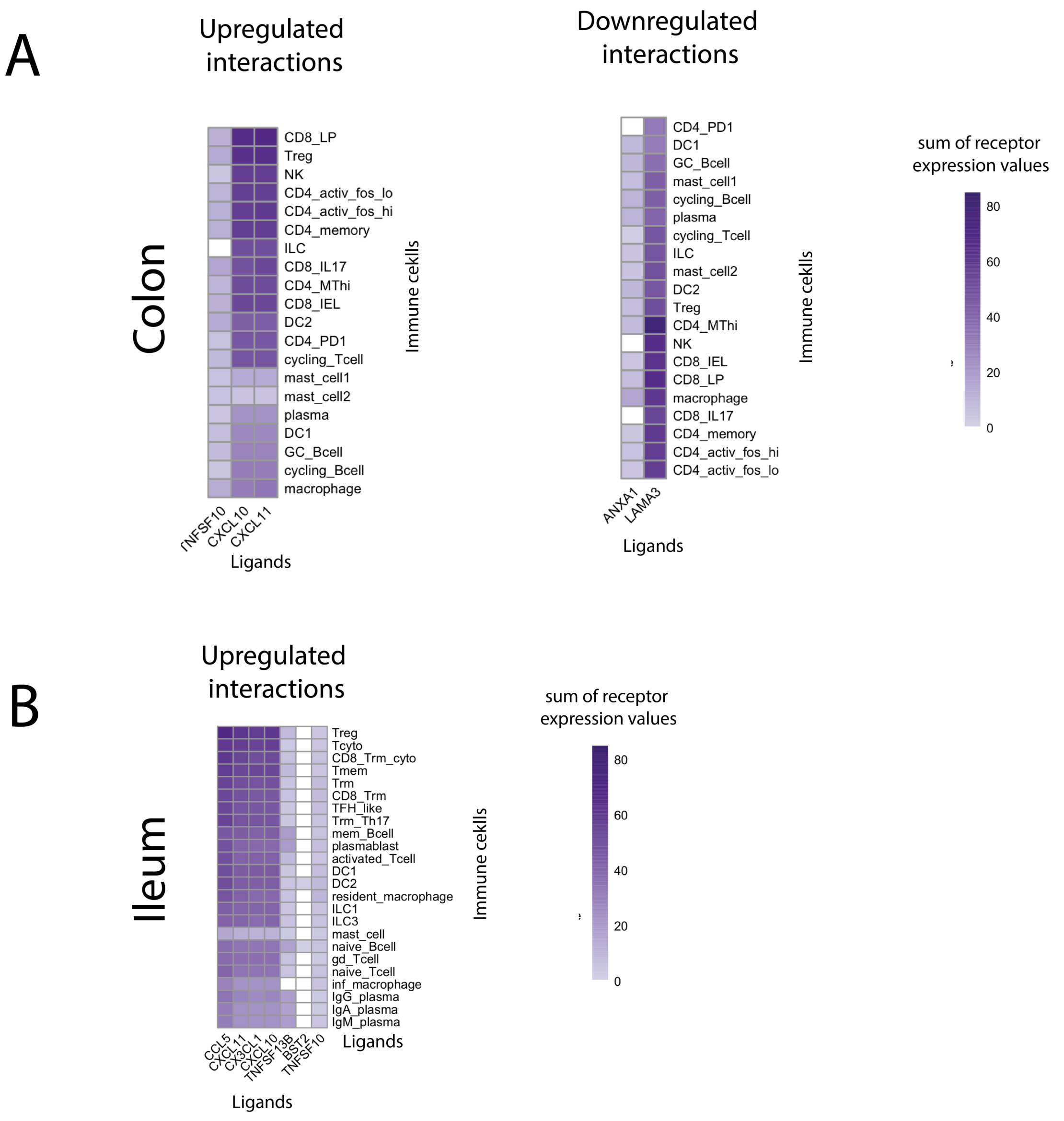


Bystander immature enterocytes
